# Sensitizing Immune-Refractory Ovarian Tumors via p53 Mutation-Tailored Immunotherapy

**DOI:** 10.1101/2025.06.23.661120

**Authors:** Rishita Chatterjee, Arturo Simoni-Nieves, An Truong, Moshit Lindzen, Furkan Ozmen, Christopher Cherry, Pascale Zwicky, Saptaparna Mukherjee, Boobash-Raj Selvadurai, Tomer-Meir Salame, Nitin Gupta, Suvendu Giri, Lior Kramarski, Yahel Avraham, Eviatar Weizman, Tugba Ozmen, Ashish Noronha, Priyasmita Chakrabarti, Deepthi Ramesh-Kumar, Julian Downward, Rony Dahan, Ido Amit, Victor Velculescu, James Brenton, Gordon Mills, Moshe Oren, Yosef Yarden

**Affiliations:** Department of Immunology and Regenerative Biology, Systems immunology, Weizmann Institute of Science, Rehovot, 76100, Israel; Cancer Research UK and Department of Oncology, University of Cambridge Cambridge, UK; Division of Oncological Sciences, Knight Cancer Institute, Oregon Health and Science University, Portland Oregon 97201 USA; Johns Hopkins University School of Medicine, Baltimore, Maryland, USA; Department of Systems Immunology, Weizmann Institute of Science, Rehovot, Israel; Francis Crick Institute, London NW1 1AT, UK; Flow Cytometry Unit, Life Sciences Core Facilities, Weizmann Institute of Science, Rehovot 76100, Israel; The Mantoux Bioinformatics institute of the Nancy and Stephen Grand Israel National Center for Personalized Medicine, Weizmann Institute of Science, Rehovot, Israel; Department of Urology, Helen Diller Family Comprehensive Cancer Center, University of California San Francisco, San Francisco, CA, USA; Department of Molecular Cell Biology, Weizmann Institute of Science, Rehovot 76100, Israel

## Abstract

High-grade serous ovarian cancer demonstrates limited responsiveness to immune checkpoint inhibitors, owing in part to immunosuppressive environments shaped by nearly universal p53 aberrations. Utilizing an immunocompetent mouse model and individual p53 mutations, we identified a dependence of the p53-R270H mutation (equivalent of human R273H) on regulatory T cells (Tregs) and the PD-1/PD-L1 axis. Analysis of patient datasets associated R273H with elevated levels of two p53 targets, PD-L1 and amphiregulin (AREG), a Tregs growth factor. In contrast to p53-R172H tumors, where there was limited activity, dual antibody therapy targeting AREG and PD-L1 selectively and effectively inhibited R270H tumors. This involved polarization toward M1 macrophages, infiltration of CD8+ T cells, diminished Ly6G+ neutrophils and downregulation of interleukin-4. In patient-derived R273C organoids, the combination treatment reduced the CD4/CD8 ratio. This study is the first to establish a mutation-tailored therapeutic approach that leverages the capacity of p53 to modulate immunosuppressive mechanisms.

## Introduction

Epithelial ovarian cancer (EOC) is considered the deadliest gynecologic malignancy [1]. Because EOC is frequently asymptomatic in its early stages, the disease is often diagnosed at advanced stages, which results in poor survival rates [2]. EOCs are categorized into Type I and Type II tumors based on histopathological heterogeneity and genetic aberrations. Type I tumors are low-grade malignancies bearing mutations in BRAF, K-RAS and PTEN, whereas Type II are high-grade tumors frequently bearing *TP53* mutations [3]. Almost 96% of patients with High-Grade Serous Ovarian Cancer (HGSOC) carry *TP53* mutations, and less than 15% of these survive 10 years post initial diagnosis [2]. Approximately 80% of the mutations are concentrated in the DNA binding domain (DBD) of this transcription factor, impairing its sequence-specific binding with DNA and resulting in loss of wild type (WT) p53 function. Some of the prevalent DBD mutations, such as R175H and R273H drive several oncogenic functions, including chemotherapy resistance [4].

Despite some advances, treatment of EOC primarily relies on surgery and chemotherapy, as well as targeted therapies, such as Olaparib, which inhibits poly (ADP-ribose) polymerase (PARP), Bevacizumab, an antibody specific to the vascular endothelial growth factor (VEGF), as well as emerging antibody drug conjugates, for example those targeting the folate receptor. PARP inhibitors are synthetic lethal with loss of *BRCA1/2* and act both as destabilizers of DNA replication forks and inducers of DNA double-strand breaks [5]. Yet, while PARP inhibitors prolong the survival of patients with *BRCA* mutations and patients with homologous recombination deficiency (HRD), resistance almost inevitably develops in PARPi-treated patients. Thus, there is a great need for better targeted treatments, especially for HGSOC. More recently, immune checkpoint inhibitors (ICIs) have emerged as an exciting new targeted therapy modality. However, unlike melanoma, lung and kidney cancer, EOC is refractory to ICIs, likely owing to low mutational burden and active immunosuppression [6].

Both tumor intrinsic and tumor extrinsic features underlie the resistance of EOC to ICIs. Poor infiltration of EOC by CD8+ effector T cells is very common [7]. Moreover, EOC frequently expresses high levels of PD-L1 to promote evasion from immune surveillance [8]. In addition to low mutational burden, which translates to relatively low abundance of neoantigens [9], EOCs frequently downregulate major histocompatibility complex class I (MHC-I) molecules, thereby preventing effective recognition by cytotoxic T lymphocytes (CTLs). Ovarian cancer cells also secrete interleukin-4 (IL-4), which directs the formation of an immunosuppressive TIME (tumor immune microenvironment) [10]. Furthermore, immune effector cells within the TIME of EOC are suppressed not only by tumor cells but also by regulatory T cells (Tregs) [11], immature dendritic cells (DCs), myeloid-derived suppressor cells (MDSCs), and cancer-associated fibroblasts (CAFs). The latter secrete immunosuppressive cytokines and chemokines (e.g., CCL2 and VEGF), which impair CD8+ T cell infiltration and activity. Together, these diverse mechanisms maintain an immunosuppressive microenvironment, permitting immune evasion [12].

Previous work showed that mutant p53 proteins can interfere with the function of the cytoplasmic DNA-sensing machinery, cGAS-STING, which activates the innate immune response [13]. In addition, the presence of missense p53 mutations negatively correlates with infiltration of CD8+ T cells into tumors of the pancreas [14], wheraes loss of p53 promotes recruitment of suppressive myeloid cells and accumulation of Tregs [15]. The proliferation of CD4+ Tregs can be enhanced in an autocrine manner by amphiregulin (AREG), a p53-regulated ligand of the epidermal growth factor receptor (EGFR) [16, 17]. We previously surveyed fluids from EOC patients and found that AREG was the most abundant and generalized EGFR ligand secreted by advanced tumors [16]. This low-affinity EGFR ligand is induced by WT p53 [18] and it undergoes upregulation following treatment with chemotherapeutic drugs [17, 18]. Interestingly, mutant forms of p53 can also upregulate AREG expression, presumably indirectly, via EGR1 [19]. Another immune-suppressive molecule regulated by p53 is programmed death ligand 1 (PD-L1) [20]. Remarkably, whereas WT p53 suppresses PD-L1 expression [21], specific mutants can upregulate PD-L1 levels [20]. In addition, unlike the translational regulation of the PD-L1 protein by mutant p53 [20], ligand-activated EGFRs stabilize PD-L1 mRNAs [22], implying that AREG and mutant p53 proteins harness different mechanisms to upregulate PD-L1.

In this study, we wished to uncover pharmacologically targetable immunosuppressive mechanisms that are activated in EOC driven by specific p53 mutants. To that end, we genetically modified a p53-deficient version of the widely used murine EOC model, ID8 [23]. Alongside ID8 cells retaining endogenous WT p53 and cells with complete p53 ablation, we stably expressed and functionally compared the murine equivalent of two human hospot p53 mutants: R270H (equivalent of human R273H) and R172H (equivalent of human R175H). Remarkably, in human EOC the R273H mutation is preferentially associated with poorer prognosis than R175H, and with upregulation of PD-L1. Consistent with the notion that R273H, but not R175H, depends on both PD-L1 and AREG, simultaneous antibody-mediated inhibition of these targets in syngeneic mouse xenografts significantly prolonged the median survival of mice carrying R270H-expressing tumors by 3.5-fold. Further analysis linked this relatively large survival benefit to increased recruitment of effector CD8+ T cells, polarization of macrophages toward the M1-like phenotype, and elevated expression of MHC-II molecules. Towards translation to clinical applications, we employed patient-derived organoids (PDO) harbouring the R273C mutation, which is very similar to R273H, and validated efficacy of the antibody combination. Together, our findings present, for the first time, a mutation-specific therapeutic strategy that exploits the nearly ubiquitous p53 mutations in HGSOC. We discuss these results in light of the continued global challenges in developing direct p53 modulators and highlight the promising alternative of targeting the distinct downstream pathways by which individual p53 mutants promote resistance to ICIs.

## Results

### p53-R273X positivity correlates with high PD-L1 and this associates with tumor aggressiveness in both EOC patients and a syngeneic animal model

Given that mutant p53 proteins engage various mechanisms to evade anti-tumor immunity—including disruption of antigen presentation and modulation of inflammatory signaling pathways [24]—we hypothesized that different p53 mutants might adopt distinct immune evasion strategies, potentially revealing mutation-specific vulnerabilities amenable to therapeutic targeting. Among the many TP53 mutations found in cancer, those affecting residues R273 (20.63%), R248 (16.67%), and R175 (14.29%) are the most prevalent [25]. Focusing on the R273? and R175? hotspot mutations, where *"?"* designates any amino acid, we conducted pan-cancer analyses of the TCGA database that revealed a consistent trend: tumors of various origins that harbored R273? mutations were associated with worse prognosis compared to those with R175? mutations (Fig. 1A). We then investigated whether this increased aggressiveness could be linked to immune evasion.

**Figure 1:**
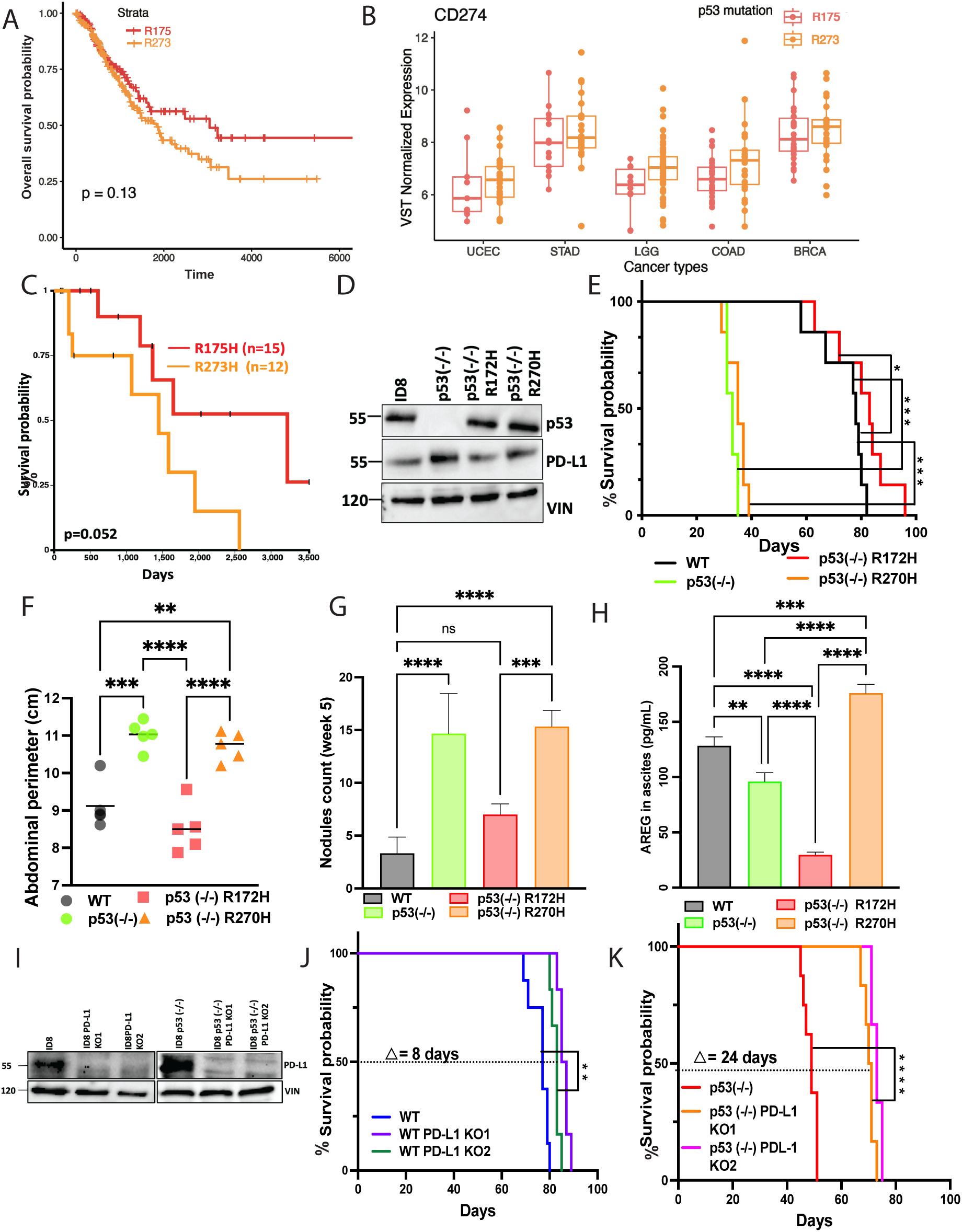
p53-R273 associates with relatively high PD-L1 and shorter survival of patients with cancer; in EOC patients and a mouse model this mutation predicts high aggressiveness relative to p53-R175. (**A**) The TCGA dataset was analyzed for survival probabilities of patients with various types of cancer harboring either p53-R175 or p53-R273 (>30 patients per cancer type). The total number of patients was 170 (R175X) and 272 (R273X). (**B**) RNAseq data from the TCGA database was used to analyze the correlation between p53 mutation status, either p53-R175X or p53-R273X, and transcript levels corresponding to PD-L1 (CD274) using variance stabilizing transformation. The cancer types analyzed were UCEC (uterine corpus endometrial carcinoma), STAD (stomach adenocarcinoma), LGG (lower grade glioma), COAD (colon adenocarcinoma) and BRCA (breast invasive carcinoma). (**C**) The TCGA-OvCA dataset was separately analyzed for survival probabilities of patients harboring either p53-R175H (n=15) or p53-R273H (n=12). (**D**) The following derivatives of ID8 cells were examined using immunoblots of whole cell extracts and the indicated antibodies: A p53-deficient stable line (p53-/-), which was used to derive two ID8 isogenic lines harboring either murine p53-R172H or murine R270H (equivalent to human R175H and R273H, respectively). (**E**) Groups of six C57/Black female mice were intraperitoneally injected with the following isogenic ID8 lines (5×10^6^ cells per animal): (i) p53(-/-), (ii) p53(-/-) cells stably expressing p53-R172H, and (iii) p53(-/-) cells stably expressing murine p53-R270H. indicated isogenic cell lines, including wildtype ID8 cells. Animal survival curves are presented. *, p< 0.05, ***, p< 0.001. (**F**) The abdominal perimeters of the mice from E were measured once per week and a representative graph for week 5 post inoculation is presented as proxy of tumor volume. (**G**) The numbers of metastatic nodules corresponding to the experimental arms shown in E were determined till week 10 post tumor cell inoculation. Shown is a graph corresponding to the fifth week’s count of nodules. (**H**) Ascites fluids were obtained from the mice of E and subjected to ELISA that measured the abundance of secreted AREG in the respective fluids. (**I**) PD-L1 knock-out clones of ID8 cells were subjected to immunoblotting to determine the protein levels of PD-L1 in the parental and PD-L1 manipulated derivatives of ID8 cells. (**J** and **K**) Groups of C57/Black female mice (n=6) were intraperitoneally injected with wildtype ID8 cells (5×10^6^ cells per animal) or derivatives lacking PD-L1 (two clones; panel J). Alternatively, mice were injected with ID8 p53(-/-) cells expressing either wildtype PD-L1 or cells whose PD-L1 was deleted (two clones; panel K). Shown are the corresponding animal survival curves. Indicated are log-rank test scores and the median gain in survival time. ****, p<0.0001.

To explore this, we identified five tumor types with at least 30 patients who carried either R273X or R175X. Using RNA-sequencing data of the corresponding tumors, we determined expression levels of PD-L1 (CD274), a well-established marker and mediator of immune escape. As shown in Figure 1B, in all five cancer types that met the selection criteria, tumors with R273X mutations exhibited higher PD-L1 expression than those with R175X mutations. These findings support the notion that p53 mutant-specific immune evasion mechanisms exist, raising the possibility that such mechanisms act in concert with cell autonomous oncogenic functions of missense p53 mutants across diverse cancer types. Focusing on high-grade EOC, we firstly confirmed shorter survival of R273H carriers relative to patients with the R175H counterpart (Fig. 1C). Hence, we next sought to address the mechanisms underlying the more aggressive and possibly more immunosuppressive phenotype of R273H in an animal model. To that end, we utilized a previously described subline of ID8 cells, a widely used mouse model of EOC. These cells express WT p53 and retain functional p53 signaling, but McNeish and colleagues used CRISPR/Cas9 gene editing to generate a p53^−/−^ derivative [26]. We transfected these p53-null cells with plasmids encoding either an empty vector or one of three forms of mouse p53: WT, R172H (mouse equivalent to human R175H) and R270H (mouse equivalent to R273H). Immunoblotting confirmed ectopic expression of the two p53 mutant proteins at similar levels (Fig. 1D). Next, all four ID8 derivatives were intraperitoneally injected into immunocompetent C57/BL6 mice to induce tumors and accumulate abdominal fluids. Animals were monitored until they reached at least one ethical end point, either a 25% increase in body weight or a 20% increase in the abdominal perimeter. The corresponding survival curves are presented in Fig. 1E. Consistent with the findings of McNeish and collaborators, mice bearing parental p53−/− tumors exhibited significantly shorter survival time compared to those with parental ID8 tumors, which retain wild-type p53. Unexpectedly, mice harboring p53-R172H tumors accumulated less ascites fluid and showed relatively prolonged survival, even slightly exceeding that of mice with WT p53 tumors (Figs. 1E and 1F). Remarkably, some mutant forms of p53 may display tumor suppressive properties in vivo, in a context-dependent manner, potentially through interference with the WNT pathway [27]. In contrast with p53-R172H, mice with p53-R270H tumors displayed very short survival durations, comparable to those with p53−/− tumors, further supporting the idea that distinct p53 mutations confer divergent effects on tumor aggressiveness. To assess the relative metastatic potential of the four ID8 isogenic lines, mice were sacrified 5 weeks post inoculation, followed by counting metastatic nodules that emerged on the inner surface of the omentum. These balloon-like lesions eventually fully covered the omental surface. Again, as shown in Fig. 1G, R270H tumors metastasized similarly to the p53^-/-^ tumors, whereas the R175H tumors were markedly less metastatic. The abundance of nodules paralleled the measured abdominal perimeters (Fig. 1F), likely reflecting the absence of physical barriers to metastasis. In line with the reported activation of the AREG gene promoter by WT p53 [18], an enzyme-linked immunosorbent assay (ELISA) demonstrated mildly greater levels of AREG in ascites fluids derived from mice carrying WT p53 tumors, relative to those carrying p53-null tumors (Fig. 1H). Remarkably, fluids isolated from mice carrying p53-R172H tumors contained significantly less AREG, whereas those associated with p53-R270H tumors displayed the highest AREG abundance, suggesting a gain-of-function effect of R270H on AREG production. Taken together, analyses of animal survival, ascites volumes, metastasis and the abundance of AREG were consistent with notable functional differences between the two missense mutations we examined.

It has recently been reported that treatment of tumor-bearing mice with recombinant AREG led to a significant reduction in intratumoral CD8^+^ T cells, along with upregulation of PD-L1 [28]. Hence, to better understand the relationships between PD-L1, p53 and AREG, we ablated using CRISPR/Cas9 the endogenous PD-L1 of both WT ID8 cells and the three p53−/− derivatives. For an unknown reason, double knockouts (p53−/−/PD-L1−/−) expressing each GOF mutant of p53 were unstable. Nevertheless, we were able to establish PD-L1-ablated clones of both WT ID8 and p53−/− cells (2 clones of each). Loss of PD-L1 was confirmed using immunoblotting (Fig. 1I) and flow cytometry (Fig. S1A). In vitro cell proliferation assays revealed that the PD-L1 knockout (KO) cells were as proliferative as the parental PD-L1-proficient cells (Fig. S1B). Nevertheless, colony formation assays uncovered involvement of PD-L1 in the ability of ID8 cells to initiate colonies in vitro (Figs. S1C and S1D). Next, we performed an in vivo experiment similar to the one presented in Figure 1E. The resulting animal survival curves are shown in Figures 1J and 1K, and photos illustrating the corresponding abdominal nodules, along with histograms of both abdominal perimeters and metastatic nodules, are presented in Fig. S1D and S1E, respectively. Ablation of PD-L1 prolonged the survival of animals harboring WT ID8 tumors by approximately 8 days, but this survival benefit was significantly prolonged to 24 days when either clone of p53-ablated ID8 cells was examined. Concordantly, PD-L1 ablation selectively reduced the number of metastatic nodules in mice carrying p53-null tumors.

In conclusion, our animal model indicates that PD-L1 is required not only for the mildly aggressive behavior of tumors with wild-type p53 but also for the heightened aggressiveness observed in p53-null tumors. These results support functional relationships between p53 and PD-L1. Importantly, in immunocompetent mice, tumors expressing p53-R172H were significantly less aggressive than those expressing R270H. Hence, our findings suggest that the frequently occurring R273H mutant of human tumors may engage preferentially both AREG and PD-L1 to promote immune evasion.

### Transcriptional profiling uncovers robust coupling of R270H, but not R172H, to redox regulation and a variant of the epithelial-to-mesenchymal transition (EMT)

In light of the different survival times displayed by xenografts of the isogenic ID8 derivatives, we performed RNA sequencing to disclose differences in the underlying mechanisms. Clustering analysis of 3-4 biological replicates of each subline (Fig. S2A) indicated low inter-sample variance. Next, we applied low-count filtering and statistical tests to list all differentially expressed genes (DEGs) in the four ID8 sublines (Fig. 2A). Notably, while depletion of WT p53 had the most pronounced impact on the transcriptome, cells expressing either point mutant differed from each other and from cells devoid of p53 (Fig. 2A). Furthermore, compared to p53 KO, R270H clearly drove greater changes in gene expression than R172H. Figure 2B presents a volcano plot showing all top DEGs differing between WT ID8 cells and cells expressing R270H. This comparison can potentially identify processes that undergo activation or repression in EOC patients when p53 acquires R273 mutations. Two important processes emerged from this analysis (note the highlighted genes in Fig. 2B): disruption of redox homeostasis and induction of EMT. The altered redox genes included several members of the glutathione S-transferase (GST) family, which catalyze the conjugation of reduced glutathione (GSH) to electrophilic substrates and detoxify oxidative stress byproducts. Among the altered genes that regulate the other major process, EMT, we identified Twist2, Ncam1 (neural cell adhesion molecule 1) and several ECM (extracellular matrix) remodelers, such as Timp3 (tissue inhibitor of metalloproteinases 3) and MMP2 (matrix metalloproteinase 2).

**Figure 2:**
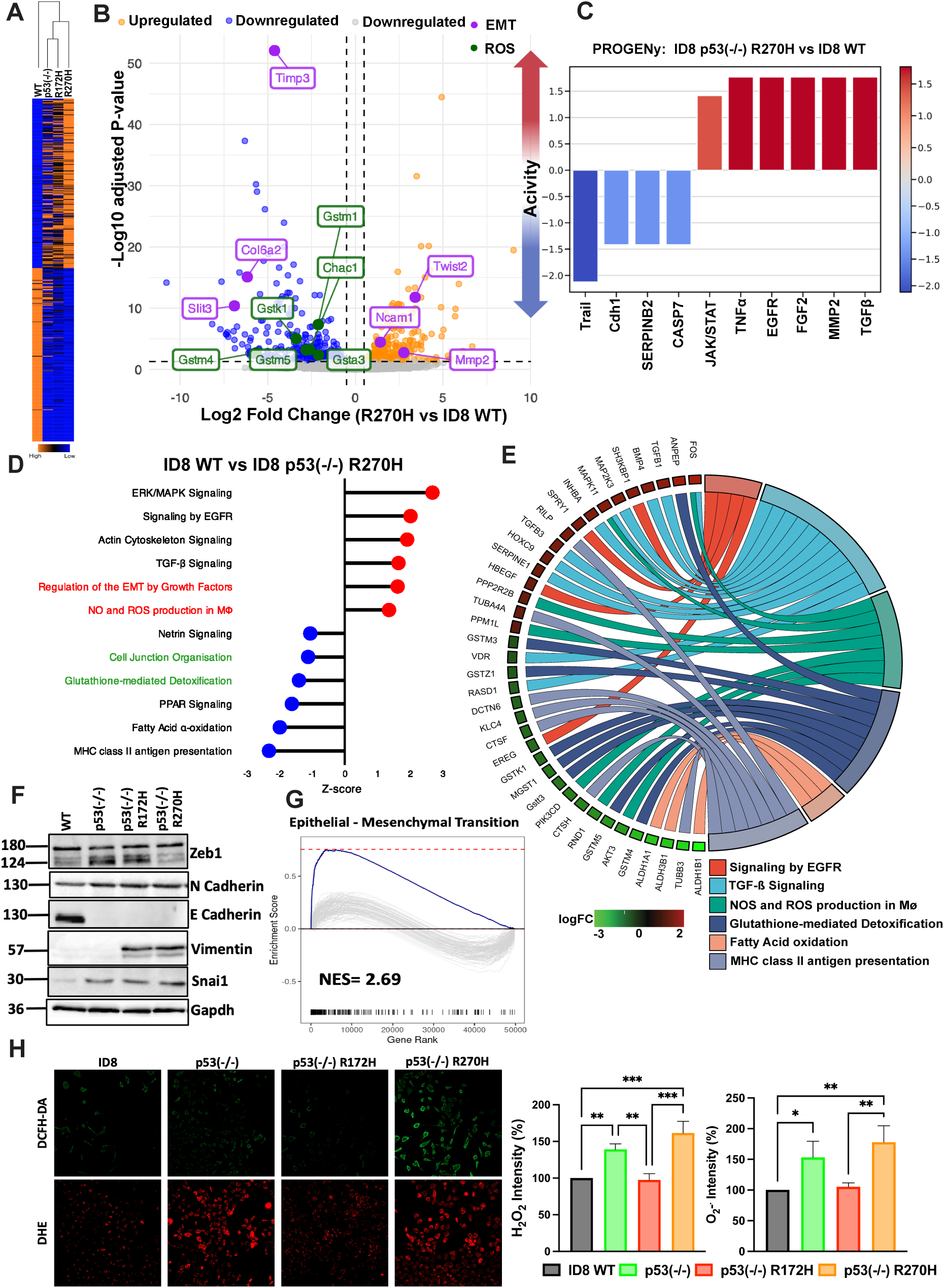
Transcriptional profiling uncovers coupling of p53-R270H, but not R172H, to redox regulation. (**A**) Bulk RNA sequencing analysis was performed on four ID8 derivatives: WT, p53-KO, p53-R172H and R270H. Shown are differentially expressed genes (DEGs) and their hierarchical clustering. (**B**) The presented Volcano plot shows all top DEGs that differed between WT and p53-R270H cells. The vertical lines mark log2 fold change of −1 and +1. (**C**) A comparative PROGENy analysis, which infers pathway activities from transcriptomics, was performed on the data shown in B. Note activation of the JAK/STAT and TGFβ signaling, as well as downregulation of Cdh1 (encoding E-cadherin) and the TNF-related apoptosis-inducing ligand (TRAIL) in ID8 p53-R270H cells. (**D** and **E**) The histogram on the left side indicates the top six canonical activated or inhibited pathways (absolute z-scores of 1 and −1, respectively) in ID8 p53-R270H as compared to wildtype ID8 cells. The right panel shows chord plots representing the same set of significant canonical pathways and displaying the relationship between individual pathways and relevant genes. (**F**) Immunoblotting analysis was performed with the indicated derivatives of ID8 cells to validate differential expression of the indicated EMT markers. GAPDH was used as the loading control. (**G)** Gene set enrichment analysis (GSEA) was applied on the DEGs of p53-R175H and p53-R273H mutated OvCa patients included in the TCGA dataset with both the KEGG pathways, as well as Hallmark gene sets. Shown is a positive enrichment score of the EMT related genes indicating that EMT related gene sets are altered at the top of the ranked list in the R273H cohort when compared to the R175H cohort. The grey lines are for individual hits, and the blue line represents the overall trend. (**H**) The indicated derivatives of ID8 cells were tested for ROS levels using either DCFH-DA or DHE, which quantify H_2_O_2_ and O_2-_, respectively. Shown are representative photomicrographs for one out of three experiments. Original magnification, 20x. Scale bar: 100-pixel size. The histograms summarize ROS quantification. *, p<0.05, **, p<0.01, ***, p<0.001.

Two different computational tools, PROGENy (Pathway RespOnsive GENes; Fig. 2C) and PEA (Pathway Enrichment Analysis; Figure 2D) confirmed that the mutation of WT p53 to R270H impacts growth factor signaling and EMT, as well as redox homeostasis. Thus, 270H is associated with higher ROS production, but lower GSH-mediated detoxification, which together predicts higher ROS accumulation. The individual proteins that pivot each process were resolved by the Chord analysis shown in Figure 2E. For example, Chord implicated *MAP2K3* and *FOS* in R270H-mediated redox regulation, likely due to redox-sensitive upstream kinases controlling MAP2K3, and ROS-sensitive FOS cysteine residues directly regulating transcriptional activity. To validate the computational findings, we immunoblotted whole extracts isolated from the four ID8 sublines. This revealed disapperance of the epithelial marker E-cadherin in the three sublines lacking WT p53 (Fig. 2F), likely due to the reported direct binding of p53 to the *CDH1* locus [34]. Correspondingly, Snail, a mesenchymal marker, was upregulated in all three sublines, but another mesenchymal marker, Vimentin, was found to be upregulated only in the two p53 mutant sublines, and a third marker, Zeb1, was upregulated in R172H but not in R270H-expressing cells. These observations indicate that loss of WT p53 induces EMT of ID8 cells and, as previously proposed [35], WT p53 and p53 mutants might exert opposing EMT effects. Still, each of the two missense mutations induced a distinct type of EMT, which might coincide with specific E/M hybrid states [36]. Importantly, our observations are consistent with previous analyses of R273H-expressing human pancreatic cancer cells [37], and they are relevant to patients with EOC because Gene Set Enrichment Analysis (GSEA) of data from EOC specimens (TCGA dataset) revealed that the EMT genesets we identified have a positive enrichment score in EOC patients expressing R273H, as compared to patients expressing R175H (Fig. 2G).

Next, we determined in vitro the content of ROS in all four isogenic ID8 sublines, using DCFH-DA (dichlorodihydrofluorescein diacetate) and DHE (dihydroethidium). Once inside cells, DCFH-DA is oxidized by ROS, such as hydrogen peroxide, to form dichlorofluorescein, which we detected primarily in R270H-expressing cells (Fig. 2H). These cells, along with the parental p53-ablated cells, also stained positively with DHE, which specifically detects superoxide (O₂.⁻). However, neither wildtype ID8 cells nor cells expressing R172H were positive for superoxide. Thus, R270H uniquely exacerbates ROS accumulation while R172H shows less ROS-centric activity. In conclusion, our ROS analyses unveiled yet another difference between the two hotspot p53 mutants. To substantiate the effects on EMT and ROS, as well as resolve the contribution of the p53-null background of the missense mutations, we performed additional pairwise comparisons:

i. ID8 p53 (-/-) versus WT cells (Figs. S2B and S2C): Applying Ingenuity Pathway Analysis (IPA) and Chord revealed activation of several oncogenic pathways in the p53^-/-^ cells, including c-MET signaling, the STAT3 pathway and catabolism of branched-chain amino acids, a source of essential nutrients required for cancer growth [29]. In line with the detection of high superoxide levels in p53^-/-^ cells, we identified glutathione-mediated detoxification as the most downregulated pathway in these cells.
ii. ID8 p53(-/-) versus R270H-expressing cells: Together, PROGENy (Fig. S2D) and PEA (Fig. S2E) uncovered activation of genes involved in the JAK/STAT and NF-κB (Nuclear Factor kappa-light-chain-enhancer of activated B cells) pathways. These routes are associated with cell survival and anti-apoptotic genes like BCL2, immune suppression and cell proliferation. Interestingly, interferon signaling was also upregulated in R270H-expressing cells, in line with the reported mutant p53-dependent STING (cyclic GMP–AMP synthase stimulator of interferon genes) activation [30].
iii. ID8 p53(-/-) versus R172H-expressing cells (Fig. S2F): The results identified downregulation of TGF-beta signaling and Nrf2 (nuclear factor erythroid 2-related factor 2), a transcription factor that normally maintains redox homeostasis. Both PTEN signaling and the Peroxisome Proliferator-Activated Receptor (PPAR) pathway were upregulated. Notably, PPARs suppress the phosphatidylinositol 3-kinase (PI3K)/Akt/Rac1 signaling axis via activation of PTEN resulting in decreased ROS, which explains the moderately low ROS levels we detected when probing R172H-expressing cells.

The relevance of these in vitro observations to patients with EOC was examined by comparing the DEGs between patients with tumors expressing R273H and those expressing R175H (TCGA dataset; Fig. S2G). The results validated differential upregulation of redox (e.g., *GSTM1*) and EMT genes (e.g., *MMP13*) in the R273H carriers. In addition, we applied GSEA on the DEGs differentiating between EOC patients harboring R273H and those harboring R175H tumors. Figure S2H presents specific EMT genes that are differentially upregulated in R273H tumors, likely due to the distinct EMT programs activated in the two sets of EOCs. Likewise, Figure S2I presents genes responsible for mitochondrial dysfunction, which were found to be downregulated in the R273H tumors. Thus, the clinical data indicate compromised mitochondrial functions in R273H tumors, relative to R175H tumors, corroborating the in vitro data. In conclusion, our analysis shows that different p53 alterations initiate mutant-specific gene programs. We also show that redox homeostasis and a variant of the EMT program are specifically activated by the relatively aggressive R270H mutation. These processes might explain the robust tumorigenic features of R270H in the ID8 animal model. Moreover, analyses of high grade human EOCs imply that the transcriptomic data extracted from our ID8 sublines bear clinical relevance for patients with advanced ovarian cancer.

### The examined mutants of p53 differentially control Nrf2 and the response to chemotherapy

To further investigate the mechanisms underlying the differential association between mutant p53 proteins and redox alterations, we firstly identified all DEGs related to redox regulation, and then analyzed their expression across the four ID8 sublines. The resulting heatmap (Fig. 3A) revealed widespread changes in antioxidant gene expression, with a particularly consistent downregulation pattern observed in the p53^-/-^ cells. Among the most prominently suppressed gene families were the glutathione S-transferases (Gsts), which detoxify ROS by conjugating them to glutathione. Specifically, transcripts for *GstA2*, *GstM1*, and *GstT1* were downregulated in both p53^-/-^ and p53 mutant cells compared to WT, while *GstA4* expression was notably elevated only in the R270H cells. Likewise *GPX4*, encoding glutathione oxidase 4, an important antioxidant that is particularly crucial in protecting against ferroptosis, was also uniquely elevated in the R270H cells. Many *Gst* genes harbor antioxidant response elements (AREs) in their promoters and are regulated by Nrf2, although some may be influenced indirectly. To assess Nrf2’s contribution, we performed RT-qPCR using primers for established Nrf2 target genes across all four cell lines (Fig. S3A). These data largely corroborated the gene expression patterns shown in Fig. 3A, supporting a central role for Nrf2 in redox regulation mediated by the status of p53.

**Figure 3:**
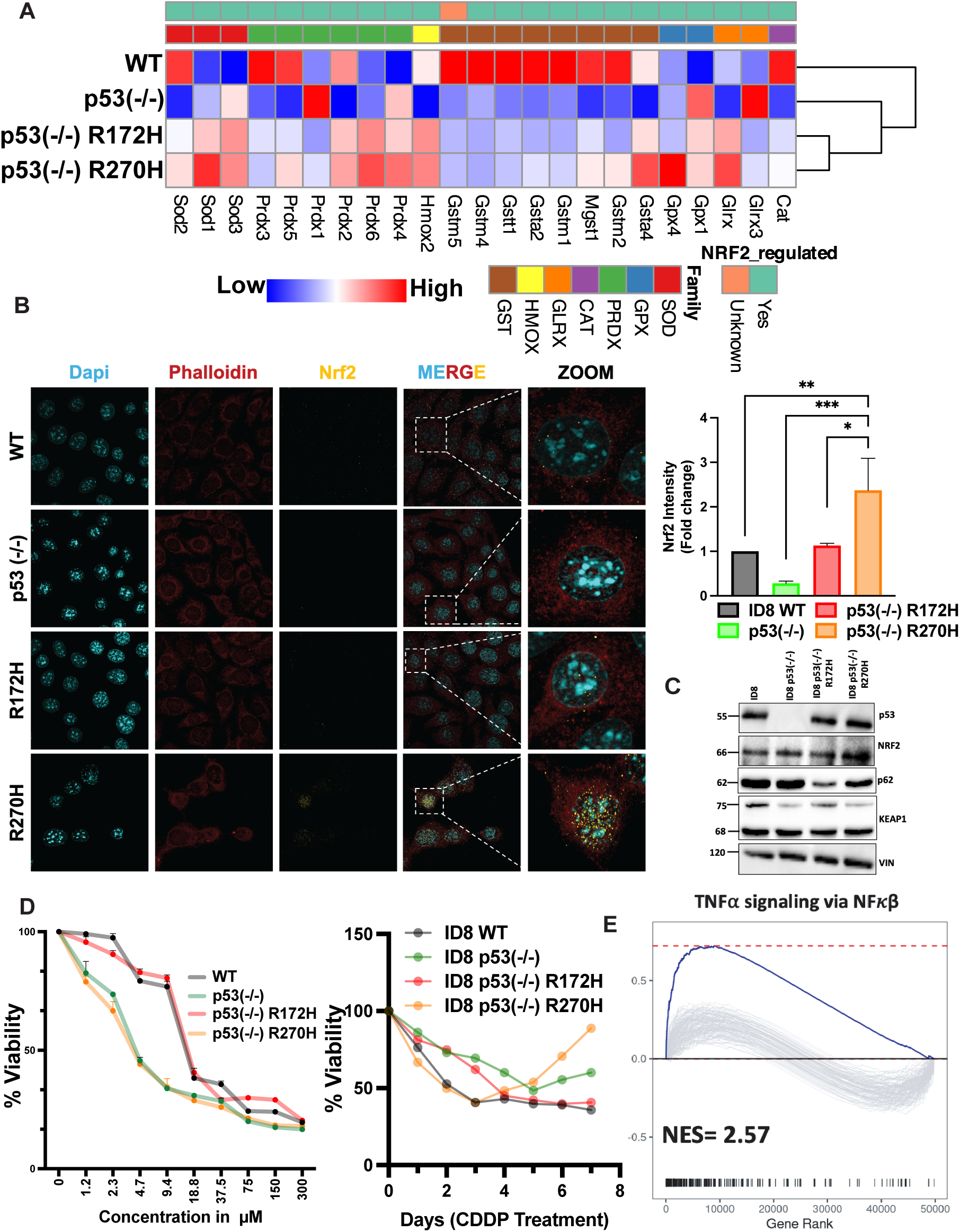
The two missense p53 mutants differentially control redox potential and Nrf2 activation. (**A**) Transcriptional profiles of redox regulatory genes displayed by the indicated ID8 sublines, including derivatives expressing p53-R270H and p53-R172H. The enzymatic antioxidant genes are functionally were seeded on coverslips and incubated overnight. Thereafter, the cells were fixed and probed using phalloidin or an anti-Nrf2 antibody, followed by a secondary antibody. DAPI was used to stain nuclei. Shown are representative images corresponding to Nrf2 immunofluorescence (green), phalloidin (red) and DAPI (blue). Bar, 20 μm. Images were captured using a confocal microscope. Nrf2 staining was quantified and normalized using ImageJ. *, p< 0.05, **, p< 0.01, ***, p<0.001. (**C**) Whole extracts of the indicated ID8 sublines were analyzed using immunoblotting for redox and autophagy markers. Vinculin was used as a gel loading control. (**D**) (left panel) The indicated ID8 derivatives (3×10^3^ cells) were seeded in 96-well plates and treated for 72 hours with increasing concentrations of Cisplatin. Cell viability was measured using the MTT (3(4,5 dimethylthiazol-2-yl)-2,5diphenyltetrazolium bromide) assay. Averages +S.D. of quadruplicates are shown. This experiment was repeated thrice. (right panel) ID8 derivatives were seeded in 96-well plates (500 cells/well) and treated with Cisplatin (3 µM). Media were refreshed once every three days. After nine days, cells were fixed and stained with crystal violet. The results of one of three experiments are shown. Note that the p53^-/-^ and the R270H lines gained resistance to CDDP. (**E**) Shown are the results of a Gene Set Enrichment Analysis (GSEA), which was applied on the differentially expressed genes between p53-R175H and p53-R273H ovarian tumors of the TCGA patient dataset (with both the KEGG pathways and the Hallmark gene sets). Note upregulation of the NF-κB pathway, mediated by TNF-alpha signaling, in the p53-R273H patients.

Since Nrf2 activity is regulated by both abundance and nuclear localization, we used confocal microscopy to assess Nrf2’s subcellular distribution. The images and corresponding quantification (Fig. 3B) showed that Nrf2 expression was reduced in p53-KO cells, whereas in R270H-expressing cells, Nrf2 was not only upregulated but also predominantly localized to the nucleus. Immunoblotting further confirmed Nrf2 upregulation in the R270H subline (Fig. 3C). In addition, the immunoblot uncovered downregulation of KEAP1, a redox sensor and substrate adaptor for the a CUL3-RBX1 E3 ubiquitin ligase complex, which ubiquitinates NRF2, targeting it for proteasomal degradation. Note the relatively low abundance of KEAP1 in p53-R270H cells, in line with their high NRF2 and reduced levels of SQSTM1/p62—a selective autophagy adaptor protein that physically interacts with p53-R273H, but not the p53-R175H mutant [31].

A key potential consequence of the differential interactions between p53 mutants and the redox system is their influence on EOC resistance to stressors, including chemotherapy. Notably, R273H has been most strongly linked to chemoresistance, for example in patients with colorectal cancer [32, 33]. To assess the relevance of these findings to EOC, we evaluated the sensitivities of the four ID8 sublines to commonly used EOC therapies: Cisplatin (CDDP), Erlotinib (an EGFR inhibitor), Auranofin (AF; a gold-based modulator of ROS), and Olaparib (a PARP inhibitor). Cell viability assays across a range of drug concentrations showed that the two ROS-enriched sublines— p53^-/-^ and R270H—were more sensitive to the ROS-targeting agents CDDP and Auranofin (Fig. S3B). In contrast, the R172H and WT ID8 cells responded similarly and did not exhibit enhanced sensitivity. We then conducted time-course experiments to assess the development of resistance. Based on the dose-response curves, we treated the sublines with CDDP (3 µM) for seven days. From Day 3 onward, CDDP-resistant clones began to emerge in the p53^-/-^ and R270H cultures (Fig. S3C), whereas no resistant clones were detected in WT or R172H cells. These findings, illustrated in Fig. 3D, demonstrate that resistance to CDDP is dependent on p53 mutation status and develops in a time- and dose-dependent manner.

To investigate the mechanisms underlying the differential resistance of the two ROS-enriched cell lines—R270H and p53^-/-^—to ROS-inducing agents such as Cisplatin (CDDP) and Auranofin, we analyzed protein expression in CDDP-treated cells via immunoblotting. As shown in Figure S3D, treatment of p53-KO and R270H cells with CDDP led to increased phosphorylation of EGFR, a response not observed in the WT or R172H sublines, which neither developed resistance nor accumulated high levels of intracellular ROS. This finding aligns with previous reports showing that R273H enhances EGFR signaling by suppressing miR-27a and that EGFR, in turn, can stabilize mutant p53 and promote drug resistance via STAT3 activation [34]. Consistently, both p53^-/-^ and R270H cells also displayed elevated NF-κB levels (Fig. 3D), supporting the established role of NF-κB in p53-related chemotherapy resistance [35]. Gene set enrichment analysis (GSEA) of EOC patient data validated the relevance of these findings (Fig. 3E). Specifically, activation of the NF-κB pathway correlated positively with the presence of the R273H mutation. Additionally, GSEA revealed upregulation of interferon-gamma signaling in patients harboring R273H (Fig. S3E), a pathway known to induce PD-L1 expression and associate with resistance to immunotherapy [36].

In summary, R270H-expressing cells, unlike those expressing R172H, exhibit elevated levels of Nrf2 and prominent nuclear localization of this transcription factor. Alongside p53-null cells, they also develop delayed resistance to treatment and maintain persistently high intracellular ROS levels. This chronically elevated oxidative stress may allow R270H cells to adapt to chemotherapy treatment through a mechanism known as “redox resetting”—a process involving Nrf2 activation and upregulation of antioxidants [37]. Through this adaptation, R270H cells may evade chemotherapy-induced cytotoxicity by activating NF-κB signaling and suppressing cell death pathways.

### Blocking either PD-L1 or AREG can prolong survival of mice bearing R270H tumors

To translate our findings into p53 mutation-specific therapeutic strategies, we first investigated the relationship between p53 status, PD-L1 expression, and prognosis of patients with EOC. We began by stratifying the WT p53 tumors in the TCGA ovarian cancer dataset into two groups, based on PD-L1 abundance: high and low. Surprisingly, higher PD-L1 expression was associated with improved overall survival in this cohort (Fig. 4A; *p* = 0.03). In contrast, among patients with p53-mutant tumors, elevated PD-L1 expression showed a trend toward worse prognosis (Fig. 4B; *p* = 0.057), suggesting oncogenic interactions between mutant p53 and PD-L1.

**Figure 4:**
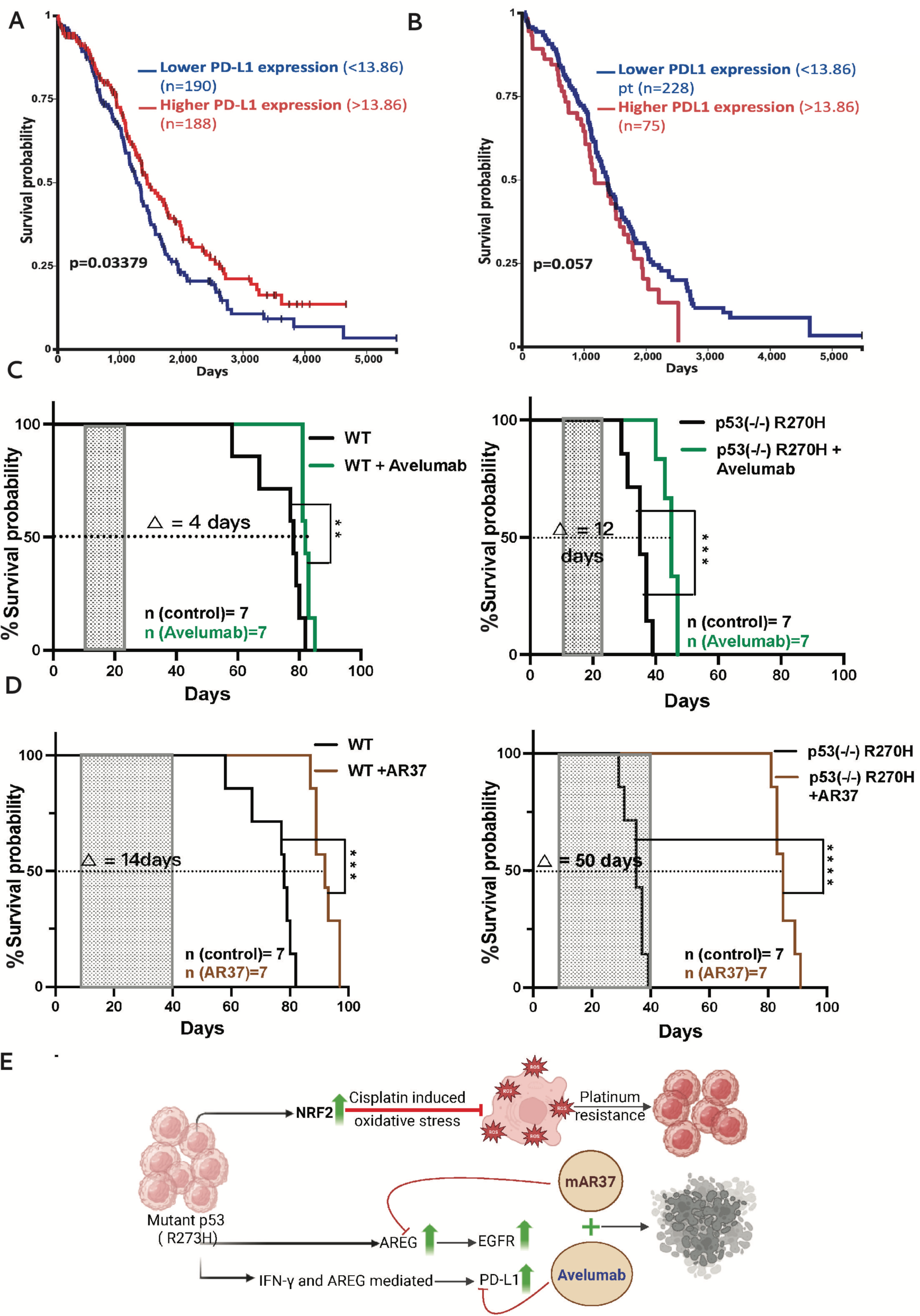
Blocking either PD-L1 or AREG extends survival of mice bearing ID8 tumors, especially tumors expressing p53-R270H. (**A**) The TCGA ovarian cancer dataset was divided into two equal groups, high and low PD-L1. Shown are the respective survival curves. (**B**) All p53-positive patients from A were divided according to PD-L1 expression levels and the respective patient survival curves are presented. Note the reciprocal trends of survival curves in A and B. (**C**) C57/Black female mice were intraperitoneally injected with 5×10^6^ ID8 cells, either WT (left panel) or p53(-/-) cells expressing a p53-R270H allele (right panel). Ten days later, mice were randomized into two groups. One group was treated with saline whereas the other received avelumab (human anti-PD-L1; 0.4 mg per injection), twice weekly starting on day ten and ending on day twenty-five. Animal survival curves are shown along with statistical analysis (**, p<0.01; ***, p<0.001). (**D**) C57/Black female mice were intraperitoneally injected with 5×10^6^ ID8 cells, either WT (left panel) or p53(-/-) cells expressing an allele of p53-R270H (right panel). Ten days later, mice were randomized into two groups. One group was treated with saline whereas the other was treated with an anti-AREG antibody (AR37; 0.2 mg/injection) from day 10 post tumor cell inoculation till day 40. Shown are the survival curves and median days gained (***, p<0.001; ****, p<0.0001). (**E**) A schematic model of the combinatorial treatment of p53-R270H ovarian cancer model using an anti-AREG antibody (AR37) in combination with an engineered anti-PD-L1 murine antibody (Avelumab).

To investigate this further in vivo, we examined how PD-L1 interacts with different p53 proteins. This set of experiments employed Avelumab, a clinically approved antibody that crossreacts with murine PD-L1. Hence, in our experiments Avelumab was used to block PD-L1 signaling in mice injected intraperitoneally with ID8 cells expressing either WT p53 or p53-R270H. Ten days post-injection, mice were randomized into control (untreated) and Avelumab-treated groups (five doses total). As expected, untreated mice bearing R270H tumors showed significantly shorter survival compared to those with WT tumors (median survival: 35 vs. 75 days). However, Avelumab treatment led to a greater survival benefit in the R270H group compared to the WT p53 group (median survival extension: 12 vs. 4 days; Fig. 4C, right panel). A parallel experiment comparing p53-KO and R172H tumors demonstrated greater aggressiveness of the p53-KO xenografts (median survival: 32 vs. 82 days), but both tumor types responded similarly to Avelumab (Fig. S4A). These results support the hypothesis that R270H, unlike R172H, functionally cooperates with PD-L1 to promote tumor progression, but this can be inhibited by Avelumab.

We next conducted a parallel series of in vivo experiments targeting AREG. For this, we used AR37, a fully murine monoclonal antibody that cross-reacts with human and mouse AREG [16, 17]. Like Avelumab, AR37 extended the survival of mice bearing WT p53 tumors (Fig. 4D; median survival gain of 14 days). However, its effect was markedly more pronounced in mice with R270H tumors, where AR37 treatment resulted in a median survival extension of 50 days (p < 0.001; Fig. 4D, right panel). In contrast, AR37 failed to significantly improve survival of mice with p53-KO or R172H tumors (Fig. S4B). Two additional outcome measures supported the anti-tumor effects of Avelumab and AR37, as well as the specificity to R270H (Fig. S4C): while neither antibody significantly reduced the number of omental nodules or the abdominal perimeter in mice with WT tumors, both agents significantly decreased these parameters in mice bearing R270H tumors. Notably, AR37 had no effect on tumors derived from p53-KO cells, consistent with the fact that these cells do not express substantial levels of AREG [17]. Taken together, findings from both animal studies and patient data support the model illustrated in Fig. 4E: R270H, unlike other p53 mutants, upregulates AREG, leading to EGFR activation, increased PD-L1 expression, and enhanced EOC progression via a pathway that engages NRF2 and antioxidant signaling.

### Combining anti-AREG and anti-PD-L1 antibodies significantly extends survival of mice bearing p53-R270H tumors, likely by enhancing antibody-dependent cellular cytotoxicity and restricting immunosuppression

To enable extended treatment regimens in tumor-bearing mice, we murinized Avelumab—originally a human antibody—by converting it to a mouse IgG2a isotype (Fig. S5A). This modification replaced the human Fc domain with a murine counterpart, while preserving the original antigen-binding region. Validation via immunoprecipitation (Fig. S5B) and ELISA (Fig. S5C) confirmed that the recombinant murine version of Avelumab retained high binding affinity for PD-L1. We then compared in mice the efficacy of this murinized Avelumab (mAvelumab) with a commercially available anti-mouse PD-L1 (mPD-L1) antibody, targeting both WT and R270H tumors. The two antibodies induced nearly identical survival benefits (Figs. S5D and S5E), confirming the functional integrity of mAvelumab and reinforcing the conclusion that PD-L1 blockade more effectively inhibits tumor progression in the R270H model (median survival: 59 days) compared to the WT p53 model (median survival: 14 days).

Since PD-L1 exerts both immunological effects through PD-1 engagement and non-immunological oncogenic functions [38], we directly compared the therapeutic outcomes of anti-PD-L1 and anti-PD-1 antibodies. Immunocompetent mice were intraperitoneally injected with either WT or R270H-expressing cells, and 10 days later they were randomized into three groups (13 mice each): untreated, anti-PD-1–treated, or anti-PD-L1–treated. Treatments continued through day 40 (10 injections total). Both antibodies yielded comparable therapeutic efficacy (Figs. 5A and 5B), with significantly stronger tumor inhibition in the R270H group relative to the WT group. These results further support the hypothesis that the aggressiveness of R270H-driven tumors depends on an active PD-1/PD-L1 signaling axis.

**Figure 5:**
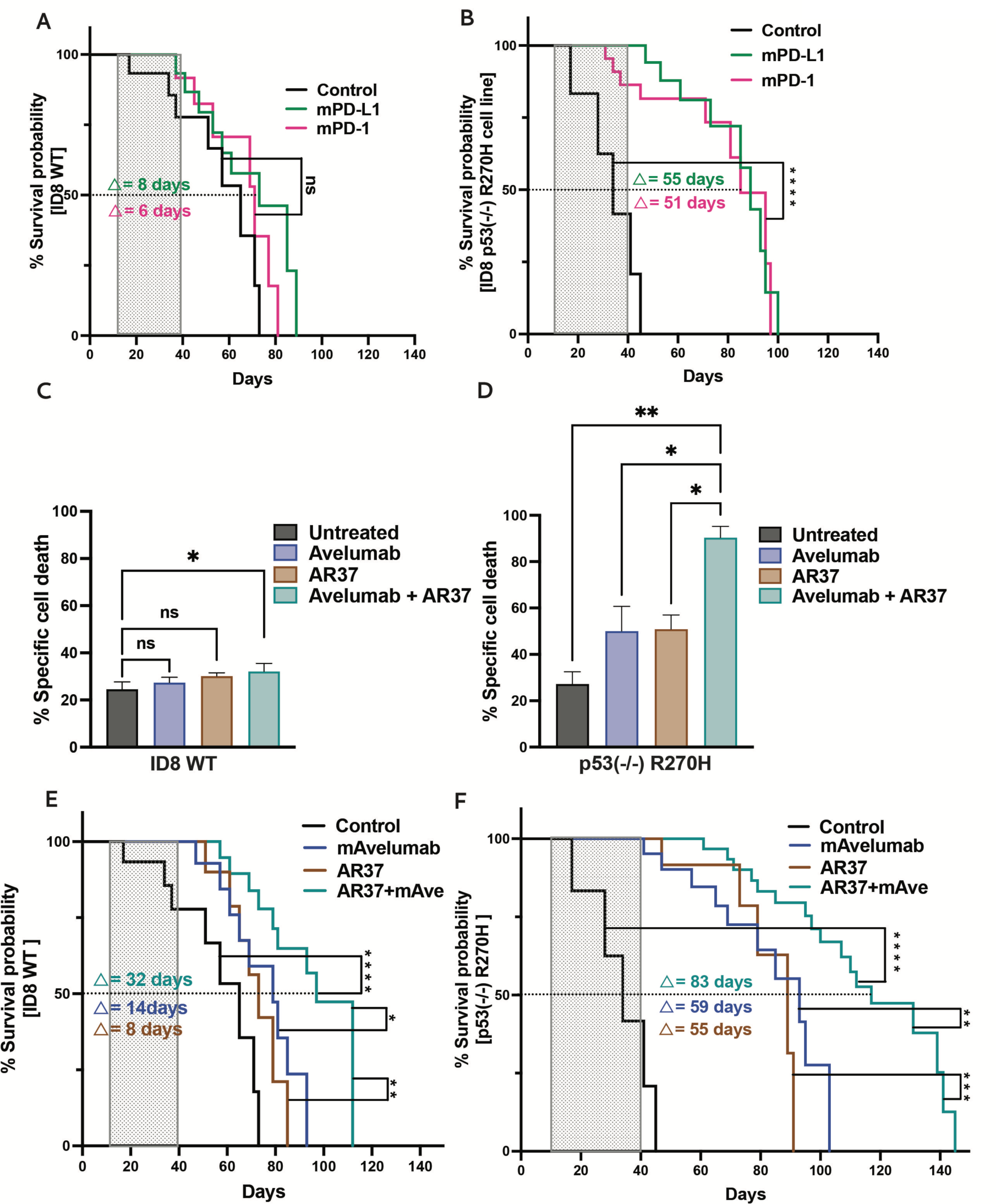
Combining anti-AREG and anti-mPD-L1 antibodies significantly prolongs survival of animals harboring p53-R270H tumors, likely by means of enhanced ADCC. (**A**) C57/Black female mice were intraperitoneally injected with p53^-/-^-ID8 cells (5×10^6^) expressing p53-R270H. Ten days later, mice were randomized into three treatment groups, each comprising 13 mice. Group 1 was left untreated, Group 2 received a commercially available mouse anti-PD-1 antibody (0.2 mg per injection) and Group 3 was intraperitoneally treated with an anti-PD-L1 antibody (from BioXcell; 0.2mg per injection). All with the median survival gains and log-rank test scores. ****, p<0.0001; n.s., not significant. (**B**) C57/Black female mice were intraperitoneally injected with p53^-/-^-ID8 cells (5×10^6^) expressing p53-R270H. Ten days later, mice were randomized into three treatment groups, each comprising 13 mice. Group 1 was left untreated, Group 2 received a commercially available mouse anti-PD-1 antibody (0.2mg per injection) and Group 3 was intraperitoneally treated with anti-PD-L1 antibody (from BioXcell: 0.2 mg per injection). All treatments terminated 30 days later (total, 10 injections). Shown are animal survival curves, along with the median survival gains and log-rank test scores. ****, p<0.0001. (**C** and **D**) Mouse peripheral blood mononuclear cells (PBMC) were isolated from bone marrows of 8 weeks old C57/Black female mice and co-cultured for 8 hours with either ID8 WT (panel C) cells or p53^-/-^ cells expressing p53-R270H (panel D), in the absence or presence of the indicated antibodies: AR37 and the engineered mouse Avelumab antibody. Note that mAbs tested in isolation were used at 30 μg/ml. When in combination, each mAb was used at 15 μg/ml. To determine ADCC efficacy, we assayed LDH release in triplicates using a kit (from Promega) that measured cell killing. The summary histogram is shown. The experiment was repeated 3 times. (**E** and **F**) C57/Black female mice were intraperitoneally injected with ID8 cells (5×10^6^), either WT or p53^-/-^ cells expressing p53-R270H. Ten days later, mice were randomized into four groups per cell line (13 mice per group). One group from each cell line was left untreated, whereas the other groups received mouse Avelumab (0.2 mg/injection), AR37 (0.2 mg/injection) or the combination of mAvelumab and AR37 (0.1 mg/injection, each). Shown are survival curves and median survival gains with a log-rank test score. *, p<0.05; **, p<0.01; ***, p< 0.001, ****, p<0.0001.

Given that mAvelumab inhibits the PD-1/PD-L1 immune checkpoint and AR37 targets the AREG-EGFR axis—crucial for regulatory T cell function [39]—we hypothesized that blocking both pathways concurrently would synergistically suppress progression of murine EOC xenografts. Prior to initiating in vivo experiments, we performed in vitro assays using co-cultures of WT or R270H-expressing ID8 cells with murine bone marrow-derived cells (BMDC), which comprised predominantly monocytes/macrophages, granulocytes, and lymphocytes. These co-cultures were maintained for 8 hours in the absence or presence of AR37, mAvelumab, or their combination. To evaluate immune-mediated cytotoxicity, release of lactate dehydrogenase (LDH) was measured. As shown in Figures 5C and 5D, ID8 cells harboring the R270H mutation were notably more vulnerable to antibody-dependent cellular cytotoxicity (ADCC) induced by either antibody compared to WT p53-expressing cells. Importantly, the antibody combination resulted in even greater killing of R270H cells than either antibody alone. Since the parental ID8 cells showed much lower ADCC responses, we concluded that the R270H mutation renders EOC cells more responsive to the dual antibody immunotherapy.

Building on the encouraging ADCC assay findings, we evaluated the therapeutic impact of the dual antibody treatment in vivo. Immunocompetent female C57BL/6 mice were intraperitoneally injected with either WT or R270H-expressing cells. Ten days later, treatments with mAvelumab, AR37, or their combination were initiated and later administered twice weekly for a total of eight doses. Following this treatment phase, tumor progression was monitored for an additional 100 days without further intervention. Mice were also routinely examined for signs of toxicity. Of note, in the combination therapy arms, each antibody was given at half the dose used in monotherapy groups. As predicted, in both tumor models the combination therapy significantly extended survival (Figs. 5E and 5F). Consistent with the in vitro ADCC data, the survival benefit was most pronounced in mice bearing R270H tumors, with median survival increasing from 35 to 115 days (a 328% improvement), compared to an extension from 63 to 100 days (59%) in the WT p53 group. Assessment of omental nodules—a surrogate for peritoneal metastases—confirmed that R270H tumors were more metastatic than their WT counterparts (Fig. S5F). Importantly, while each antibody alone reduced nodule burden, their combination proved more effective.

In summary, dual targeting of PD-L1 and AREG resulted in superior therapeutic benefit compared to monotherapy, particularly in tumors harboring the R270H mutation. This enhanced efficacy likely reflects the combined effect of promoting anti-tumor immunity while simultaneously suppressing tumor-supportive immune cell populations.

### Co-cultures comprising ID8 derivatives, bone marrow cells and patient-derived organoids validate that simultaneous inhibition of PD-L1 and AREG increases CD8-to-CD4 ratios

Patient-derived organoids (PDO) have demonstrated particular promise as novel ex vivo systems that faithfully recapitulate the response of EOC to anti-cancer drugs [40]. To ensure stable antibody recognition of the human forms of AREG and PD-L1, we used Surface Plasmon Resonance (SPR), along with recombinant forms of both antigens. The sensorgrams shown in Figs. S6A and S6B display SPR responses reflecting high affinity binding of the chimerized form of AR37 (comprising a human Fc) toward human AREG and of Avelumab toward hPD-L1.

Next, three models of HGSOC were selected from a PDO repository [41]. All three models carry p53 mutations, either C176F (PDO1; Fig. 6A) or R273C (PDO5 and PDO6). To simulate physiological T cell activation prior to co-culturing with matched peripheral blood mononuclear cells (PBMC, derived from the corresponding EOC patient), we first stimulated the PBMC in vitro using CD3/CD28-coated beads (Dynabeads™ Human T-Activator CD3/CD28, Gibco). These beads mimic antigen-presenting cell signals by engaging CD3, a component of the T-cell receptor, and CD28, a co-stimulatory molecule. PBMC were activated for 48 hours before initiating co-cultures. The activated cells were then subjected to treatment with human Avelumab, a partly humanized version of AR37 (bearing a human Fc), or a combination of both antibodies.

**Figure 6:**
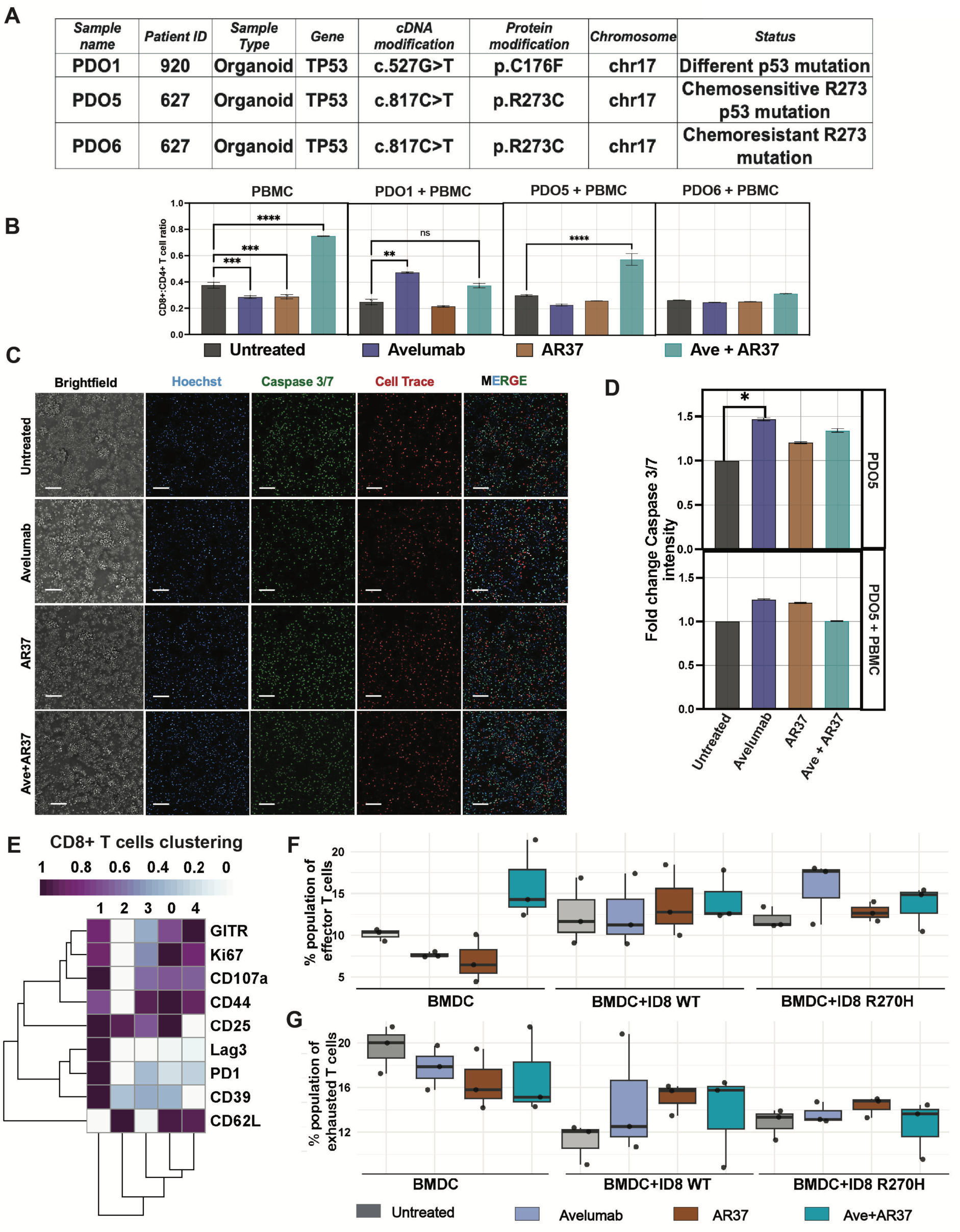
Treatment of patient-derived OvCA organoids with a combination of anti PD-L1 and anti-AREG antibodies reveals extensive CD8+ T cell recruitment, which is confirmed using a murine co-culture model. (**A**) Tabular description of the PDO models selected for the coculture experiment. (**B**) Cytometry-based estimation of the ratio between CD8+ and CD4+ T cells in co-cultures. Patient-matched PBMC were incubated for 5 days with three different PDOs. The incubations were performed in the presence of indicated antibodies. After the fifth day, the organoid-PBMC cultures were dissociated and subjected to flow cytometry with antibodies specific for CD8+ and CD4+, and the event count for each of these cell types was taken to calculate the ratio. The PBMC only used healthy PBMC. Note that PDO5 and PDO6 express p53-R273C. (**C**) Shown are results of an imaging-based apoptosis assay depending on detection of caspase-3/7 activation and a fluorogenic substrate. The indicated PDOs were co-cultured with human PBMC (3×10^5^ cells) for 5 days and treated with specific antibodies (15 micrograms per millilitre). (**D**) Quantification of the caspase activation signals only from the PDO cells is provided in histograms. Note that Avelumab displayed an apoptotic effect toward the chemo-sensitive PDO5. Statistical analysis used two-way ANOVA with a log-rank test score. **, p<0.01, ***, p<0.001 and ****, p<0.0001 is shown. Scale bar, 100 µm. (**E**) Mouse bone marrow derived cells (BMDCs) were co-cultured for 2 hours with ID8 isogenic cell lines, either wildtype or p53(-/-) R270H cells. Afterwards, cells were stained for surface or nuclear antigens to identify the following T cell subtypes using flow cytometry: naïve and effector memory (cluster 0), exhausted (cluster 1), early differentiated (cluster 2), memory (cluster 3) and effector T cells (cluster 4). (**F** and **G**) Boxplots showing the fractions of exhausted CD8+ T cells (F; cluster 1) and effector CD8+ T cells (G; cluster 4) across the indicated untreated or antibody-treated BMDC, which were cultured either alone or together with murine ID8 cells, either wildtype or p53(-/-) R270H cells.

Two functional assays were employed: one assayed apoptotic markers (as previously described [42]), and the other, a flow cytometry-based assay, that determined the CD8-to-CD4 T cell ratio. The latter revealed a marked increase in CD8+ T cells relative to CD4+ T cells following a 72-hour long treatment of PBMC monocultures from a healthy donor with the antibody combination (Fig. 6B). These results suggest that the dual antibody treatment effectively boosts cytotoxic T lymphocyte populations while potentially reducing the proportion of regulatory CD4+ T cells [43]. Co-culturing PDO5 (R273C; chemosensitive) with the cognate patient’s PBMC supported the ability of the antibodies to AREG and PD-L1, when combined, to increase the CD8/CD4 ratio (Fig. 6B). However, neither antibody was active alone and, furthermore, neither PDO6 (R273C; chemoresistant) nor PDO1 (p53-C176F), replicated the superiority of the antibody combination. Note, however, that PDO1 did respond to Avelumab monotherapy. These observations exemplify the complexity of patient-derived PDO models and the matched PBMC cocultures. Specifically, it is well known that PBMC’s lymphocytes and myeloid cells are short lived and they display high sensitivity to prior patient treatment with chemotherapy. Considering the caveat that an antibody targeting AREG, a soluble rather than a surface-bound antigen, may not trigger robust immune cell death, we assessed programmed cell death by measuring activation of caspase-3 and caspase-7. The monotherapies, as well as the antibody combination, were monitored in all three PDO co-cultures and their respective patient-derived PBMC. Four-channel imaging was used to estimate the apoptotic signals corresponding to the organoid cells only, while filtering out signals emanating from the PBMC. The results obtained with PDO5 (R273C) showed the ability of Avelumab monotherapy to induce apoptosis in the monoculture setting (Figs. 6C and 6D). However, this was not enhanced by the inclusion of the patient’s PBMC, nor by adding antibody AR37. Similar results were obtained when analysing PDO6 (R273C, chemoresistant; Figs. S6C and S6D), which responded to Avelumab in the absence of PBMC, and PDO1 (C176F; Fig. S6D), which responded to Avelumab in the presence of PBMC.

To corroborate the findings from the PDO-PBMC co-cultures, we applied an antibody panel on co-cultures comprising ID8 derivative cell lines and mouse BMDC. The panel included CD8+ T cell markers of differentiation and degranulation (CD107a), proliferation (Ki67), migration/homing (CD62L), effector functions (GITR) and T cell exhaustion (LAG3, PD-1 and CD39) [44–46]. Mouse bone marrow was derived from three healthy immunocompetent mice and processed for isolating BMDC, which were then incubated overnight. Co-cultures were established with the ID8 WT and the p53-R270H cell lines, which were treated with antibodies having mouse Fc region, with the goal of capturing Fc-mediated immunomodulatory functions. A lineage clustering was performed to identify CD8+ T cells subtypes across all samples. Agreeably, the combination of AR37 and Avelumab increased the CD8+ T cell population in triplet co-cultures (Fig S6E). Even though the impact of AR37 monotherapy on CD8-positive cells was lower, it exceeded the effect Avelumab monotherapy. To identify the involved subtypes of CD8+ T cells, we used the aforementioned panel of markers and segregated all T cells into the main subtypes (Figs. 6E and S6F). We were especially interested in T cell exhaustion, hence focused on cluster 1 (exhausted T cells) and cluster 4 (effector T cells). This analysis uncovered a tendency of the antibody combination to elevate the fraction of cytotoxic T lymphocytes (CTLs), alongside a stronger tendency of the combined treatment to diminish the fraction of exhausted T cells (Figs. 6F and 6G), the major drivers of immunosuppression within the TIME. This in vitro observation prompted us to further explore the immunomodulatory effect of the combination therapy in the immunocompetent mouse model.

In conclusion, analyses of PDOs supported the observations made when studying an animal model of EOC. Specifically, mutations affecting residue Arg273 of human p53 enabled a chemosensitive PDO model to increase the TIME’s CD8/CD4 ratio in response to treatment with a combination of antibodies blocking both AREG and PD-L1. In addition, an antibody blocking PD-L1 significantly increased death of the respective cancer cells, but inclusion of the anti-AREG antibody did not significantly enhance cancer cell death in this assay. Similar to the results obtained with ovarian PDOs and PBMC, co-culture experiments using ID8 sublines and BMDC concluded that the combined antibody treatment can reduce the exhausted subtype of CD8+ T cells and concurrently increase cytotoxic T cells. This bimodal mechanism likely underlies the observed inhibitory impact of the pair of antibodies on animal survival.

### Concurrently blocking AREG and PD-L1 in R270H-expressing tumors polarizes macrophages, recruits CD8+ T cells and diminishes both CD38+ myeloid cells and Ly6G+ neutrophils

CyTOF (Cytometry by Time of Flight) can simultaneously measure multiple parameters at the single-cell level, which permits a comprehensive characterization of the TIME [47]. A preselected panel of metal-conjugated anti-mouse antibodies was used to probe ascites fluids for recruitment of specific mouse immune cells to the TIME. To resolve mechanisms underlying the response of the R270H tumors to immunotherapy, we treated tumor-bearing mice with our murine anti-AREG and anti-PD-L1 antibodies. All treatments started on day 10 and ascites fluids were collected 65 days later. In addition to CyTOF, we applied cytokine arrays and analyzed metastatic nodules (see a schematic flow chart in Fig. S7A).

Following treatment of mice with anti-AREG and anti-PD-L1 antibodies, CyTOF detected changes in several markers in the collected ascites fluids. These included MHC-II, PD-L1 (CD274), CD38, CD8 and F4/80. Figure 7A lists all markers in a dot plot that displays single-cell expression data and the corresponding immune cell clusters, such as tumor associated neutrophils (TANs). Figure 7B shows a viSNE plot of CD274 (PD-L1), utilizing a visualization tool together with the t-SNE (t-distributed stochastic neighbor embedding) algorithm, which demonstrated downregulation of PD-L1 following 35 days of treatment of R270H tumors with the combination of AR37 and Avelumab. Note that Avelumab’s IgG1 backbone enables it to recruit natural killer (NK) cells to lyse PD-L1–expressing tumor cells via ADCC [48]. Manual gating for the CD3-positive population in the ascites fluid, representing T cells, established yet another feature of the antibody combination, namely enhanced recruitment of the CD8 positive fraction of T cells (Figs. 7C and S7B).

**Figure 7:**
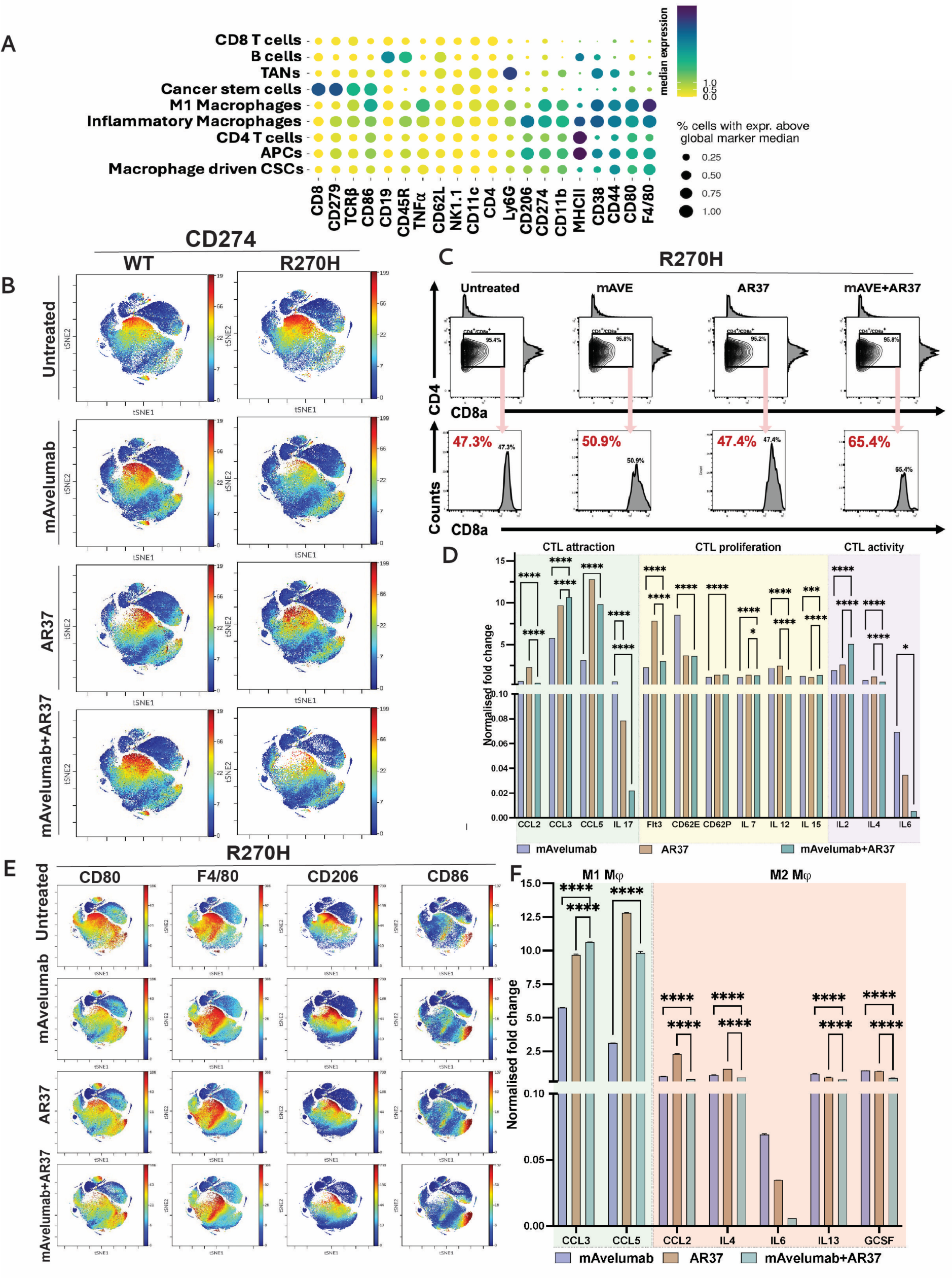
Concurrent blockade of AREG and PD-L1 drives recruitment of CD8^+^ T cells and M1 macrophage to p53-R270H tumors. (**A**) C57/Black female mice intraperitoneally injected with ID8 cells (5×10^6^), either WT or p53^-/-^ cells overexpressing p53-R273H. Ten days later, mice were randomized into four groups per cell line (5 mice per group). One group was left untreated, whereas the other groups received treatment with either mAvelumab (0.2 mg/injection), AR37 (0.2 mg/injection) or a combination of mAvelumab and AR37 (0.1 mg/injection, each). Ascites fluids were collected on day 75 post cell inoculation and processed for CyTOF analysis. The presented dot plot shows the differential association of the preselected immune cell markers with specific cell populations that were present in the ascites fluid of untreated mice bearing p53-R270H tumors or mice treated with the antibody combination. Dot size refers to the fraction of cells with marker expression above global marker median, and dot color marks the median abundance of the respective marker. Note the variation in markers of T cells, macrophages and antigen presenting cells. (**B**) viSNE plots of the results of CyTOF analysis of single cells in mouse ascites fluids using an antibody specific to CD274 (PD-L1). Note that wildtype ID8 cells are compared to p53-R270H in terms of the cellular response to two antibodies, AR37 (anti-AREG) and mAvelumab, as well as the combination of the two antibodies. The coloration is proportional to the expression intensity (red = high). Note downregulation of the PD-L1 cell population in the antibody combination group. (**C**) Shown are analyses of CD3+ T cells in ascites fluids. This analysis used manual gating. Note increased CD8-positive T cells following treatment of the p53-R270H cells with the combination of two antibodies. (**D**) Ascites fluids from two mice were used to overlay cytokine array membranes (from Proteome Profiler^TM^ Mouse Cytokine Arrays). Duplicate spots corresponding to 111 cytokines were assayed and the obtained data were normalized to the untreated group. Note antibody-induced changes in the abundance of cytokines involved in CTL recruitment (CCL2, CCL3, CCL5 and IL-17), CTL proliferation (Flt3 ligand, E-selectin, P-selectin, IL7, IL12 and IL15) and CTL activation (IL2, IL4 and IL6). (**E**) viSNE plots corresponding to the following macrophage markers: CD80, F4/80, CD206 and CD86. Note diminished abundance of M2 macrophage markers and concurrent upregulation of M1 markers upon treatment of the p53-R270H model. (**F**) Ascites fluids from two mice were used to overlay cytokine array membranes as in D. The obtained data were normalized to the untreated group. Note changes in the abundance of cytokines responsible for M1-like macrophage polarization (left colored box; CCL3 and CCL5), which were upregulated, along with downregulation of M2-promoting cytokines (right colored panel; CCL2, IL4, IL6, IL13 and GCSF). ****, p<0.0001.

The viSNE plot and the data shown in Figures S7C and S7D illustrate yet another feature of the simultaneous blockade of AREG and PD-L1 in R270H-expressing mice: downregulation of both CD38+ myeloid cells and Ly6G+ neutrophils. Notably, TANs inhibit cytotoxic T lymphocytes by depleting L-arginine and releasing ROS. Neutrophils also induce T-cell exhaustion and suppression of Th1, NK and B cells [49]. The other therapy-induced marker we detected, CD38, is a multifunctional molecule expressed on plasma cells, T and NK cells. The noted decrease in CD38+ myeloid cells suggests that simultaneously blocking AREG and PD-L1 delivers a therapeutic effect by inhibiting CD38-mediated T cell exhaustion [50].

Next, we analyzed the cytokines accumulated in the peritoneum of mice bearing WT or R270H tumors, prior to and post treatment with the antibody doublet. Following the isolation of fluids from two animals per treatment arm, these fluids were overlaid on cytokine array membranes (Proteome Profiler^TM^ Mouse Cytokine Arrays). Duplicate spots corresponding to 111 cytokines were probed and the resulting data were normalized against the untreated group. The results are presented in Figure 7D. Notably, the levels of multiple cytokines playing roles in CTL recruitment (CCL2, CCL3, CCL5 and IL-17), proliferation (Flt3 ligand, E-selectin, P-selectin, IL-7, IL-12 and IL-15) and activation (IL-2, IL-4 and IL-6) were altered. Remarkably, IL-4 was strongly downregulated post treatment with either anti-PD-L1 or the antibody combination. This finding is consistent with a recent report demonstrating that ovarian cancer cells secrete IL-4, which directs the formation of an immunosuppressive TIME [10]. Likewise, IL-2 underwent upregulation in response to the dual antibody treatment, likely reflecting its ability to expand cytotoxic T and NK cells. Similarly, the levels of chemokine ligand 2 (CCL2) and other cytokines were altered in a manner that is similarly consistent with their roles in the recruitment of suppressive monocytes and macrophages to the ovarian TIME [51]. The dot plot shown in Figure S7E classifies the altered cytokines into two categories: (i) immunomodulatory cytokines, such as interferons and TNF-alpha (tumor necrosis factor alpha), and (ii) compensatory growth factors, such as EGF and CXCL1, which activate EGFR either directly or indirectly [17]. Of note, this and other pathways were variably influenced by each antibody, but all were inhibited by the dual antibody combination, consistent with the high anti-tumor efficacy of this treatment strategy.

In summary, our in vivo EOC models displayed multiple molecular alterations. Yet, three features were commonly observed: (i) in comparison to the application of each antibody alone, the combination of anti-AREG and anti-PD-L1 antibodies elicited clearly stronger effects on Ly6G+ neutrophils, PD-L1, and cytokines such as IL-4, (ii) relative to mice carrying WT ID8 tumors, the effects displayed by mice carrying R270H tumors were typically stronger, and (iii) all observed changes were consistent with attenuation of the immunosuppressive nature of the TIME.

### Combining anti-AREG and anti-PD-L1 antibodies increases MHC-II and induces polarization of macrophages towards M1

The differentiation of macrophages to a pro-inflammatory M1-like phenotype or an anti-inflammatory M2-like phenotype plays critical roles in shaping the TIME. Using CyTOF analysis and viSNE plots (Fig. 7E), we noted that the antibody combination diminished in mice carrying R270H tumors several markers associated with M2-like macrophages (e.g., CD206) and enhanced markers associated with M1-like macrophages, such as CD80 and CD86. In addition, we noted an increase in the abundance of the major histocompatibility complex type II (MHC-II) (Fig. S7F), implying enhanced antigen presentation and immunomodulatory effects. Concordantly, the use of cytokine arrays confirmed that factors responsible for M1-like macrophage polarization, such as CCL3 and CCL5, were upregulated following treatment with Avelumab and AR-37, either as monotherapies or in combination (Fig. 7F). Reciprocally, the combinatorial treatment induced the downregulation of several M2-associated cytokines, such as G-CSF [52] and additional factors (e.g., CCL2, IL-4, IL-6 and IL-13).

In summary, this study investigated *TP53*—the most commonly mutated gene in epithelial ovarian cancer (EOC)—with a focus on the functional diversity of its mutations. The central hypothesis was that different p53 mutants, due to their unique interacting proteins and effects on transcriptional programs, may drive immunosuppression and resistance to ICIs via distinct mechanisms. Clinical data guided the focus on the R273H mutation, which appeared to rely on two immune-related factors: regulatory T cell (Treg)-mediated immune suppression and a functional PD-1/PD-L1 pathway. Using an immunocompetent mouse model, we demonstrated that the murine equivalent, R270H, unlike R172H or complete p53 ablation, requires both amphiregulin—a Treg-associated growth factor—and PD-1/PD-L1 signaling. As predicted, dual blockade of these independent pathways with specific antibodies suppressed tumor growth through multiple mechanisms, including effector T cell activation, M1-like macrophage polarization, and modulation of cytokine profiles. These results suggest a new therapeutic strategy for a particularly aggressive EOC subtype and support mutation-specific immunotherapeutic approaches capable of transforming immune “cold” ovarian tumors into inflamed, treatment-responsive ones.

## Discussion

Currently, only a limited number of isogenic murine cell models of EOC suitable for xenograft studies in immunocompetent mice are available. One recently reported model was developed using murine fallopian tube epithelial cells with combined loss of *p53*, *Brca1*, *Pten*, and *Nf1*, along with overexpression of *Myc* and *p53*-*R172H* [53]. Similarly, another study created mouse fallopian tube organoids to establish homologous recombination-proficient models lacking *p53* [54]. Since these models were not yet available when our study began, we utilized a *p53*-deficient derivative of the widely used, spontaneously immortalized ID8 ovarian cancer cell line [26]. Utilizing ID8 cells, we previously identified AREG as the most abundant and broadly expressed EGFR ligand in ascites fluids from advanced EOC patients [16, 17]. Therefore, we generated a neutralizing anti-AREG antibody (in AREG-deficient mice), which cross-reacts with murine and human AREG [54]. This antibody inhibited the growth of ID8 xenografts and significantly enhanced the effectiveness of chemotherapy. The underlying mechanism likely involves inhibition of a pathway initiated by p53 binding to the AREG promoter and culminating in activation of EGFR signaling [17]. Parallel studies have shown that AREG promotes the immunosuppressive function of regulatory T cells (Tregs) [39]. These findings led us to pursue a dual-targeting approach, inhibiting both AREG and the PD-1/PD-L1 axis. This was motivated by the low response rates of human EOC to ICIs (<10%) [55] and the reported ability of high AREG levels to predict response of colorectal cancer patients to anti-EGFR antibodies [56]. Our results unraveled an unanticipated feature of p53: in comparison to tumors expressing WT p53, no p53 or p53-R172H, treatments combining anti-AREG and anti-PD-L1 antibodies were significantly superior in prolonging the survival (by >3-fold) of animals carrying tumors expressing the R270H mutant.

To appreciate the notable efficacy and mutation-specific action of the dual antibody therapy we describe herein, it is useful to compare targeting mutant forms of p53 and targeting the RAS family— the second most frequently mutated protein in tumors. Although p53 mutations are even more prevalent, no p53-based therapies have so far been approved by the Food and Drug Administration. This stands in contrast to the RAS pathway, where inhibitors such as Sotorasib and Adagrasib have recently received clinical approval, and several others are progressing through late-stage trials. These RAS inhibitors act by binding directly to the effector-binding domain and other functional regions of RAS proteins. Mutant p53 proteins, however, present a greater challenge due to the absence of binding pockets for small molecules, with the exception of Y220C that is presently being targeted in clinical trials [57]. Consequently, rather than targeting mutant p53 itself, our approach focused on inhibiting two critical downstream effectors—AREG and PD-L1. Importantly, both targets are involved in the ability of mutant p53 to mediate immune evasion [39, 58], and similar to other EGFR ligands [22], AREG independently elevates PD-L1. In addition, human R273X mutations are particularly frequent not only in EOC but also in cancers of the brain, pancreas, and liver, where this residue has the highest mutation frequency [59]. R273H is noted for its ability to increase genomic instability, boosts resistance to certain chemotherapies, as well as mediates distinct protein-protein interactions [32, 60, 61]. Conceivably, because R273H modulates specific immunosuppression mechanisms, blocking these routes might expose previously unrecognized vulnerabilities.

When effective, immunotherapy typically induces several parallel changes that collectively remodel the TIME to favor anti-tumor immunity. Employing in vivo models, cultured cells and PDOs, we validated that concurrent inhibition of both AREG and PD-L1 in R270H-expressing cells changed both the repertoire of immune cells and the landscape of TIME’s cytokines. Firstly, elevated levels of CD8+ T cells were detected and, in parallel, CD4-positive T cells were depleted from the ascites fluid of treated mice. Similarly, we observed several indications for polarization of macrophages towards an M1-like state, implicated in production of pro-inflammatory cytokines and ROS, which can kill tumor cells [62]. M1-like macrophages also promote Th1 responses, which are crucial for CTL recruitment and activation. Yet another alteration we detected was the depletion of CD38+ myeloid cells. CD38 is an ectoenzyme that converts NAD+ to adenosine, thereby leading to adenosine accumulation, which suppresses anti-tumor immunity. Notably, depleting CD38+ cells reverses T cell exhaustion and enhances cytotoxicity. Similarly, we observed depletion of Lys6G+ neutrophils in the ascites fluids of treated mice. This alteration typically relieves suppression of CD8+ T and NK cells, leading to more effective cytotoxic responses [63]. In conjunction with elevated IL2, which can enhance the cytotoxic activity of NK and T cells, and decreased IL4, which associates with an immunosuppressive TIME [10], these alterations boost anti-tumor immunity and reverse T cell exhaustion.

Given the multifaceted effects on the TIME, the high therapeutic efficacy observed in animal models, and the specificity to the R270H mutation, our findings warrant serious consideration for translational development. Notably, despite the approval of targeted therapies such as Bevacizumab and PARP inhibitors, the 5-year survival rate for EOC has shown little improvement over the past decade, and the disease remains highly lethal [64]. Contrary to initial expectations, EOC has proven largely unresponsive to ICIs, due in part to low tumor mutational burden and an immunosuppressive TIME. Apart from hereditary BRCA1 and BRCA2 germline mutations, most somatic alterations—such as in PIK3CA or ARID1A—are confined to specific EOC subtypes. In contrast, TP53 mutations are almost universal in HGSOC, with R273 variants (e.g., R273H and R273C) comprising 20.6% of TP53 hotspot mutations in this subtype [65]. Furthermore, our approach targets a defined patient group for whom chemotherapy currently remains the only option. The combination of antibodies against AREG and PD-L1 may offer a tolerable safety profile: the main adverse effects of anti-PD-L1 therapies involve immune-related reactions, such as diarrhea and skin inflammation [66]. While no approved therapies currently target AREG, studies show that AREG-deficient mice are viable and fertile, with their most pronounced phenotype being impaired mammary gland development during puberty [67]. Future studies will assess the comparative efficacy of combining anti-AREG antibodies with either anti-PD-1 or anti-PD-L1 antibodies. Additionally, it will be critical to evaluate whether omental AREG and tumor PD-L1 expression levels can serve as predictive biomarkers for the proposed immunotherapy regimen.

In summary, it is well established that the cell-autonomous effects of mutant p53 proteins are complemented by their ability to promote evasion from both innate and adaptive immune responses. Our findings indicate that distinct p53 mutations engage different signaling pathways and immunosuppressive mechanisms. Hence, by uncovering p53 mutation-specific processes of immune evasion it may be possible to bypass the need for direct p53 targeting—as exemplified in this study through dual antibody blockade of AREG and PD-L1. This antibody combination extensively alters the TIME, hence demonstrates strong efficacy in immunocompetent mice. Given the near-universal presence of p53 mutations in HGSOC and the limited treatment options beyond chemotherapy, our approach offers a promising therapeutic path and a foundation for future mutation-tailored strategies targeting p53.

## Acknowledgements

We thank all members of our laboratory for their insightful comments. This work was performed in the Marvin Tanner Laboratory for Research on Cancer. YY is the incumbent of the Harold and Zelda Goldenberg Professorial Chair in Molecular Cell Biology. Funding: This work was supported by the Israel Science Foundation, the European Research Council (ERC), the Israel Cancer Research Fund (ICRF), the National Cancer Institute (U01 CA 281902, to GBM) and the Dr. Miriam and Sheldon G. Adelson Medical Research Foundation (AMRF).

## Data and materials availability

Transcriptomics data have been submitted to Gene Expression Omnibus (GEO), a public functional genomics data repository (https://www.ncbi.nlm.nih.gov/gds) under the code GSE283235. All materials that are not commercially available will be made available to interested readers. Please contact the lead author, Dr. Yosef Yarden.

## Materials and methods

### Cell lines and reagents

The ID8 wildtype and the p53^-/-^ derivative were kindly provided by Prof. Iain McNeish (Imperial College, London). Two derivatives were established in our lab: ID8 p53^-/-^ ectopically expressing murine p53-R172H, which represents the human equivalent, R175H, and ID8 p53^-/-^ cells expressing p53 R270H (representing the human equivalent, R273H). In addition, we established ID8 derivatives lacking PD-L1. All ID8 sublines were cultured in Dulbecco’s modified Eagle’s (DME) medium supplemented with FBS (10%), penicillin, streptomycin and sodium pyruvate. Mouse PBMC were isolated from mouse bone marrow and grown in Roswell Park Memorial Institute (RPMI) medium supplemented with 10% FBS. Plasmid transfections used jetPEI.

### Generation of stable cell lines

Knockout of PD-L1 in wildtype ID8 cells and in ID8 p53^-/-^ cells was made using the CRISPR-Cas9 system. This involved the creation of a double-stranded break next to the Protospacer Adjacent Motif (PAM). The target site was selected using the ENSEMBL database and the selected 21 bp target included the PAM sequence in exon 3.

### Cell growth and colony formation assays

To assay cell viability, we employed the MTT (methoxynitrosulfophenyl-tetrazolium carboxanilide) assay. ID8 cells (3×10^3^) were seeded in 96-well plates in quadruplicates and treated with drugs for 72 hours. Next, MTT was added, and plates were incubated for three hours at 37°C. Optical density was measured at 490 nm and 640 nm. For colony formation assays, cells were seeded in 6-well plates and monitored for 14 days, replenishing the media once every three days. On days 2, 4, 6, 8, 10, 12 and 14, cells were fixed in cold methanol and stained with crystal violet. Images were captured and analyzed using ImageJ. Later, crystal violet was dissolved using 2% SDS, and absorbance was measured at 590 nm.

### Nucleotide sequencing of RNA

RNA isolation was performed using Dynabeads mRNA Direct Kit (from Thermo Fischer Scientific). NGS libraries were prepared using a modified version of Transeq, where RNA was barcoded and reverse-transcribed using poly-T primers, followed by the addition of an exonuclease to remove excess PCR primers. The single-stranded cDNA was then converted to double-stranded DNA. Template DNA was removed using DNase, and the RNA generated was fragmented and ligated to barcoded Illumina adapters. Reverse transcription of the ligated product was performed using primers specific for Illumina’s adapters, and cDNA libraries were generated and enriched by performing 12-15 PCR cycles. RNA-seq libraries (pooled at equimolar concentrations) were sequenced on an Illumina NextSeq 500 at a median sequencing depth of 10 million reads per sample. Sequences were mapped to the mouse genome, demultiplexed and filtered.

### RNA-seq data analyses and pathway analyses

Poly-A/T stretches and Illumina adapters were trimmed from the reads using cutadapt, and reads shorter than 30bp were removed. The filtered reads were mapped to the mouse reference genome GRCm38 mm10. Reads with identical Unique Molecular Identifiers (UMIs) were removed using the PICARD MarkDuplicate tool. Gene expression levels were quantified using htseq-count. Differentially expressed genes (DEGs) were identified using DESeq2 with betaPrior, cooksCutoff and independent filtering parameters, which were set to False. Initial p-values were adjusted for multiple testing using the Benjamini and Hochberg procedure. The analysis pipeline was managed using Sneakmake. Once a group-wise comparison was made, the DEGs were subjected to core analysis that used the Ingenuity Pathway Analysis (IPA) platform. Selected pathways with valid z-scores were chosen with fold change of DEGs of at least 1 or −1, and an adjusted p-value smaller than 0.05.

### RNA isolation and real-time PCR analyses

Total RNA extraction was performed using the PerfectPure RNA Cell Kit (5prime, Hamburg) following the manufacturer’s instructions. RNA quantity and quality were determined using the Nanodrop ND-1000 spectrophotometer (Thermo Fischer Scientific). Complementary DNA strand was synthesized using the High-Capacity Reverse Transcription kit (Applied Biosystems). Real-time qPCR analysis was performed with SYBR Green system (Applied Biosystems) and specific primers on the StepOnePlus Real-Time PCR system (Applied Biosystems). qPCR signals (CT) values were normalized to beta2microglobulin (B2M).

### Cell lysis and immunoblotting

As mentioned in the figure, post-treatment cells were washed using ice-cold saline. Whole cell lysates were collected in a mild lysis buffer (50 mM HEPES, pH 7.5, 10% glycerol, 150 mM NaCl 1% Triton X-100, 1 mM EDTA, 1 mM EGTA, 10 mM NaF and 30 mM β-glycerol phosphate). Cell lysates were normalized by estimating protein content using BCA and then immunoblotted. Membranes were blocked with TBS-T (tris-buffered saline containing Tween-20) containing 1% low-fat milk, blotted overnight with a primary antibody, washed thrice in TBS-T, incubated for 1 hour with a secondary antibody linked to horseradish peroxidase, and rewashed three times with TBS-T. Immunoreactive bands were detected using the ECL reagent (Biorad)

### ROS determination assays

ID8 derivative lines were seeded in 12-well plates and on the following day, hydrogen peroxide and anion superoxide were assessed by incubating cells with DCFDA (10 μM) and DHE (10 μM, diluted in Krebs-Ringer phosphate buffer) for 30 minutes in the dark at 37°C and under 5% CO_2_. After washing the cells twice in Ringer phosphate buffer, cellular fluorescence was measured using epifluorescence microscopy (Olympus Corporation, Japan) at an excitation wavelength of 485/500 nm (DFCDA/DHE) and an emission wavelength of 530/580 nm (DFCDA/DHE). Signals were quantified using ImageJ.

### In vivo animal experiments

All animal experiments were approved by the Weizmann Institute’s Animal Care and Use Committee and the Institute’s Review Board (IRB). C57/Black female mice of 5-6 weeks of age were intraperitoneally injected with ID8 cells (5 x 10^6^), either WT, p53^-/-^, p53^-/-^ R172H and p53^-/-^ R270H. Next, animals were randomized into the indicated treatment groups, such as the commercially available (from BioXcell) murine anti-PD-1, or anti-PD-L1 (intraperitoneal injections). Treatments started on day 10 post cancer cell inoculation and lasted until day 40 (for antibodies) or day 45 (for other drugs). The measured abdominal perimeters, along with animal survival curves, are shown along with a log-rank test score. Upon euthanasia, ascites fluid, ovaries, fallopian tubes and metastatic nodules were collected for further analysis.

### Cloning and production of murine Avelumab

The sequences encoding the heavy and light chains of Avelumab were used for the cloning. A double digestion with Age-I and Not-1 was performed for the heavy and light chains, respectively. PCR cleanup was performed after digestion, followed by a ligation step that used the digested vector backbone plasmid DNA for sequencing. Approximately 8 µl of DNA, considering the plasmid’s DNA concentration, was used for DNA sequencing. At least 1 µg of DNA was diluted in water (18 µl), and 2 µl of CMV primers were next added and used for sequencing. Only positive clones were selected and the plasmids were amplified using the NucleoBond Xtra Midi Kit to obtain transfection-grade plasmids. To produce mAvelumab, plasmids encoding the heavy and light chains were transfected using ExpiFectamine into Expi293 cells grown in suspension. To boost production, enhancers 1 and 2 were added on the next day, as per the Expi293 kit instructions. Following a week-long incubation, the antibody was purified using a Protein G column and later underwent dialysis against saline.

### Surface plasmon resonance (SPR) measurements

Binding affinity and dissociation of the antigen-antibody complexes were tested by surface plasmon resonance using the BIAcore 200 instrument. PD-L1 and AREG proteins obtained from Sino-Biological were tested for purity first and then immobilized on CM5 chips in different channels. HBS-EP (10 mM HEPES with 0.15 M NaCl, 3 mM EDTA, and 0.05% surfactant P20, pH 7.4) was used as the running buffer. An increasing concentration of the antibodies Avelumab (0.01 to 5 nM) and AR37 (0.02 to 10 nM) were passed through the channels immobilized with respective antigens. The dissociation constant Kd and association constant Ka were calculated. Sensograms were fitted to a steady-state model (T200 software). All experiments involved two independent repeats.

### Immunoprecipitation assays

Protein A/G agarose beads were coated with human or mouse Avelumab for 1 hour at 4°C, on a rotor. Next, the beads were washed and incubated at 4°C, overnight, with a recombinant mouse PD-L1 extracellular domain (His-tagged, from Vector Builders). On the next day, the beads were thoroughly washed and the antigen eluted in concentrated gel loading buffer to validate the pull-down efficacy of the engineered antibodies.

### Antibody-dependent cell-mediated cytotoxicity (ADCC) assays

Mouse PBMC were isolated from the bone marrow of 8-week-old C57/Black female mice. The marrow of the femur bone was thoroughly suspended in ice-cold saline and treated with ACK (Ammonium-Chloride-Potassium) lysis buffer to remove red blood cells. Next, bone marrow cells were subjected to MARS CS cell separation and co-cultured for 8 hours with ID8 cells or with their derivatives, without or with therapeutic antibodies (Avelumab 30 μg/ml, AR37 30 μg/ml and the combination arm; 15 μg/ml of each antibody) to capture the ADCC efficacy of the mouse Fc region shared by the two antibodies. As a measure of cell death, we assayed LDH release using a kit from Promega.

### Establishment of patient-derived ovarian cancer organoids

Three previously described organoid models (PDO1, PDO5 and PDO6) [41] were suspended in 0.02 ml domes consisting of 80% BME (Cultrex Reduced Growth Factor Basement Membrane Extract Type 2, from Bio-Techne R&D Systems) and 20% ADF++ media, consisting of Advanced DME/F-12 medium (Gibco) supplemented with 12 mM HEPES and 2 mM glutamine. Organoid domes were grown in GyneCult™ Fallopian Tube Organoid Medium (FT media, from Stem Cell Technologies) consisting of ADF++ media, Wnt3a-conditioned medium (25% v/v), RSPO1-conditioned medium (25% v/v; in-house), 1X B27 (an optimized serum-free supplement; ThermoFisher), N-2, a chemically-defined, serum-free supplement (1x, from ThermoFisher), recombinant Noggin protein (100 ng/ml; Peprotech), human epidermal growth factor (10 ng/ml; Peprotech), nicotinamide (1 mM; Sigma), 0.5 μM SB431542 (an ALK5 inhibitor, from Cambridge Biosciences), 9 μM Y27632 (RHO kinase inhibitor, from Cambridge Bioscience), 100 Units/ml penicillin and 0.1 mg/ml streptomycin (Gibco).

### PBMC cultures and activation

PBMC were obtained frozen from BPS Biosciences. All experiments were performed with PBMC from the same batch (Cat no. 79059-1, Lot no. 230808). PBMC were thawed three days prior to co-culture set up for T-cell activation. Frozen PBMC were thawed into RPMI 1640 media supplemented with 10% heat-inactivated FBS, 2 mM sodium pyruvate, 100 Units/ml penicillin and 100 μg/ml streptomycin (Gibco) and counted using a manual haemocytometer. Dynabeads™ Human T-Activator CD3/CD28 for T Cell Expansion and Activation (Gibco) were added at a ratio of 25 μl per 10^6^ PBMC. After three days, cell suspensions were placed on a magnet to remove the Dynabeads.

### BMDC isolation

Bone marrow derived cells were isolated from healthy C57/Black female mice of 6 weeks age by humanly sacrificing the animals followed by extraction of bone marrow from the femur and tibia bones. The marrow was thoroughly suspended in ice-cold saline and treated with ACK (Ammonium-Chloride-Potassium) lysis buffer to remove red blood cells. Next, bone marrow cells were subjected to MARS CS cell separation and cultured for 8 hours prior to establishing the co-cultures with ID8 isogenic cells.

### Organoids and ID8 co-cultures

Organoids were dissociated from BME domes using TrypLE (Gibco) and resuspended in FT media. PBMC separated from Dynabeads were also resuspended in FT media. Next, organoids and PBMC were mixed at 1:20 ratio and suspended in FT media with 10% BME. Before plating, culture plates were pre-coated with an anti-human CD3 monoclonal antibody (1 μg/ml; Invitrogen) and with anti-human CD28 mAb (3 μg/ml; Invitrogen). For fluorescence microscopy, organoid (cancer) cells were stained with Hoechst 33342 (0.5 μg/ml), and PBMC, stained with CellTrace Far Red (ThermoFisher) and washed twice with saline prior to mixing. For apoptosis assays, we co-incubated 5000 organoid cells and 100,000 PBMC in 0.05 ml suspension containing 10% BME. The cell suspension was dispensed in 96-well black optical plates (ThermoFisher). Each well received 0.05 ml FT media containing CellEvent Caspase-3/7 dye (ThermoFisher), which was added after BME polymerization. For flow cytometry, cells were not stained prior to mixing. 0.05 ml of the co-culture suspensions (in 10% BME) containing 15,000 organoid cells and 300,000 PBMC were dispensed in 48-well culture plates (Corning), and FT media (0.15 ml) containing Cell Event Caspase-3/7 dye (ThermoFisher), which was added after BME polymerization. Three organoid models were selected from the repository of several PDOs based on the p53 status. ID8 isogenic cells lines (WT and p53^-/-^ R270H) were grown and counted and seeded in 24 well plate with the isolated BMDC cells at a ratio of 1:20, without or with therapeutic antibodies (Avelumab, 30 μg/ml; AR37, 30 μg/ml and the combination arm; 15 μg/ml of each antibody) for two hours. Co-cultures were then processed for staining of surface and intranuclear markers for FACS using eBioscience™ Foxp3 / Transcription Factor Staining Buffer Set.

### Fluorescence microscopy and image analysis

Cells were imaged using the Operetta CLS™ high-content analysis system (Revvity). Images were obtained using a non-confocal, water objective (20x) across 4 channels. Imaging fields across eight z-stacks at a step size of 5 μm were obtained per well. Maximum projections across z-stacks were generated for each channel using the BIOP plugin in FIJI (https://github.com/BIOP). An analysis pipeline was constructed using Cell Profiler. Organoid nuclei were segmented using Hoechst 33342 intensities. Any regions overlapping with PBMC, as segmented using CellTrace Far Red intensities, were discarded. CellEvent caspase-3/7 dye intensity was then measured for non-overlapping nuclear regions, effectively quantifying levels of apoptosis in organoid cells.

### Flow cytometry analyses

Prior to flow cytometry, co-cultures were dissociated with TrypLE and resuspended in buffer consisting of saline supplemented with FBS (10%). Cells were stained for 1 hour prior to fixation with PFA (4%). Staining made use of Hoechst 33342 (0.5 μg/ml) and an antibody mixture consisting of anti-human APC-EpCAM, PE/Cy7-CD3, BV750-CD8 and BV605-CD4 (all antibodies were from BioLegend). Next, cells were analyzed using a flow cytometer (the BD LSRFortessa™). Organoid cells were gated using the epithelial marker EpCAM, and T-cells were gated using CD3. The CD3^+^ population was subdivided into a CD4^+^ T-helper and CD8^+^ cytotoxic T-cell populations. CellEvent Caspase-3/7 (Invitrogen) intensities were measured to quantify apoptosis. Prior to flow cytometry for the ID8-BMDC cocultures using the FACS Aurora, co-cultures were transferred in PBS to 96 well v-bottomed plate. Cells were washed in 0.2 ml PBS per well. 50ul of surface staining Mastermix was added to the cells with 50ul of “live/dead only” was added to the control and unstained well had 50ul of PBS. Cells were then incubated in the dark at 4 degrees for 30 minutes. Fixation was done after washing the cells and cells were later incubated at 4 degrees for 30 minutes. A wash with Wash buffer/ perm buffer was done to prepare the cells for intracellular staining followed by overnight intracellular staining antibodies. All the samples including the unstained and live/dead stained samples were washed in 0.15 ml of PBS. Cell samples were stored in 0.1 ml of PBS, at 4 degrees in the dark until the acquisition using the FACS Aurora after filtering out the clumped cells, if any. The data acquired was unmixed and then gating was performed to analyze the live BMDC alone to capture the changes in T cells sub population upon the antibody treatment.

### CyTOF analyses of ascites fluids

Cells collected from fresh ascites fluid from antibody-treated and control animals harboring ovarian cancer were subjected to ACK lysis to eliminate red blood cells. Following a brief DNase treatment, cells were incubated with Fc-Receptor blocking solution from BD Biosciences (Mouse TruStain FcX PLUS #156603/156604; 5 μl per 10^6^ cells). Cells were washed thoroughly and subjected to dead cell staining. Samples were CD45 barcoded after being suspended in the Maxpar cell staining buffer. Surface staining was performed using a cocktail of metal-conjugated selected antibodies characterized by different metal masses. Samples were stained with conjugated Cisplatin (194Pt/195Pt/196Pt/198Pt) at 1.25 µM, then deactivated using a complete RPMI medium and consecutive washing steps. Final fixation of the pooled samples was performed using 4% freshly made PFA in Maxpar PBS (in ampules; from Pierce; 16% Formaldehyde (w/v), methanol-free. Cells were further stained with Cell ID Intercalator-Ir-125 µM (Iridium solution at 250 nM in 4% PFA). Each sample was washed thoroughly and suspended in Maxpar Cell Acquisition Solution Plus (CAS+). For bead preparation and data acquisition we applied the following protocol: 1:10 CyTOF EQ Four Element Calibration beads (EQ Beads) were resuspended in CAS+ solution. Cells in the sample were counted and gravity filtration was done three times to remove clumps and debris using a 35 µm filter mesh cell strainer (Falcon). A final solution of cells (2×10^6^/ml) and beads was set up in the CyTOF-Helios, which used a cytometer (Fluidigm). The data was pre-processed to normalize and concatenate prior to analysis by Cytobank. Data was normalized using CyTOF Software (v.6.7.1014). Gates were applied using the Cytobank platform (Beckman Coulter). The normalized beads were gated out, and then live and dead cells were gated using Cisplatin (194Pt) and Iridium DNA label (193Ir). Event length and Gaussian parameters were used considering the width, center, offset and residual channels. CyTOF software was then used for sample de-barcoding. The de-barcoded data set was analyzed using FlowJo to count the dead cell population, followed by CytoBank analysis to create the tSNE plots.

### Analysis of ascites fluids using cytokine arrays

Following the removal of cells, fresh ascites fluids isolated from antibody-treated mice were overlaid on cytokine array membranes (Proteome Profiler^TM^ Mouse Cytokine Arrays). The membranes were developed using streptavidin-HRP and signals were measured using the Chemidoc BioRad system. The measured duplicate spots corresponding to 111 cytokines were analyzed using Image Studio software by LiCOR Systems. Data was normalized to the untreated group.

### Statistical analyses

Data were analyzed using the Prism Graphpad software and R. Statistical analyses were performed using t-test and one- or two-way ANOVA with Tukey’s or Dunnett’s tests and log rank tests (*, p<0.05; **, p<0.01; ***, p<0.001; ****, p<0.0001). All experiments were repeated at least thrice.

