## Supplementary Figures for "Sensitizing Immune-Refractory Ovarian Tumors via p53 Mutation-Tailored Immunotherapy"

**Rishita Chatterjee et al. Supplementary Figures and legends**

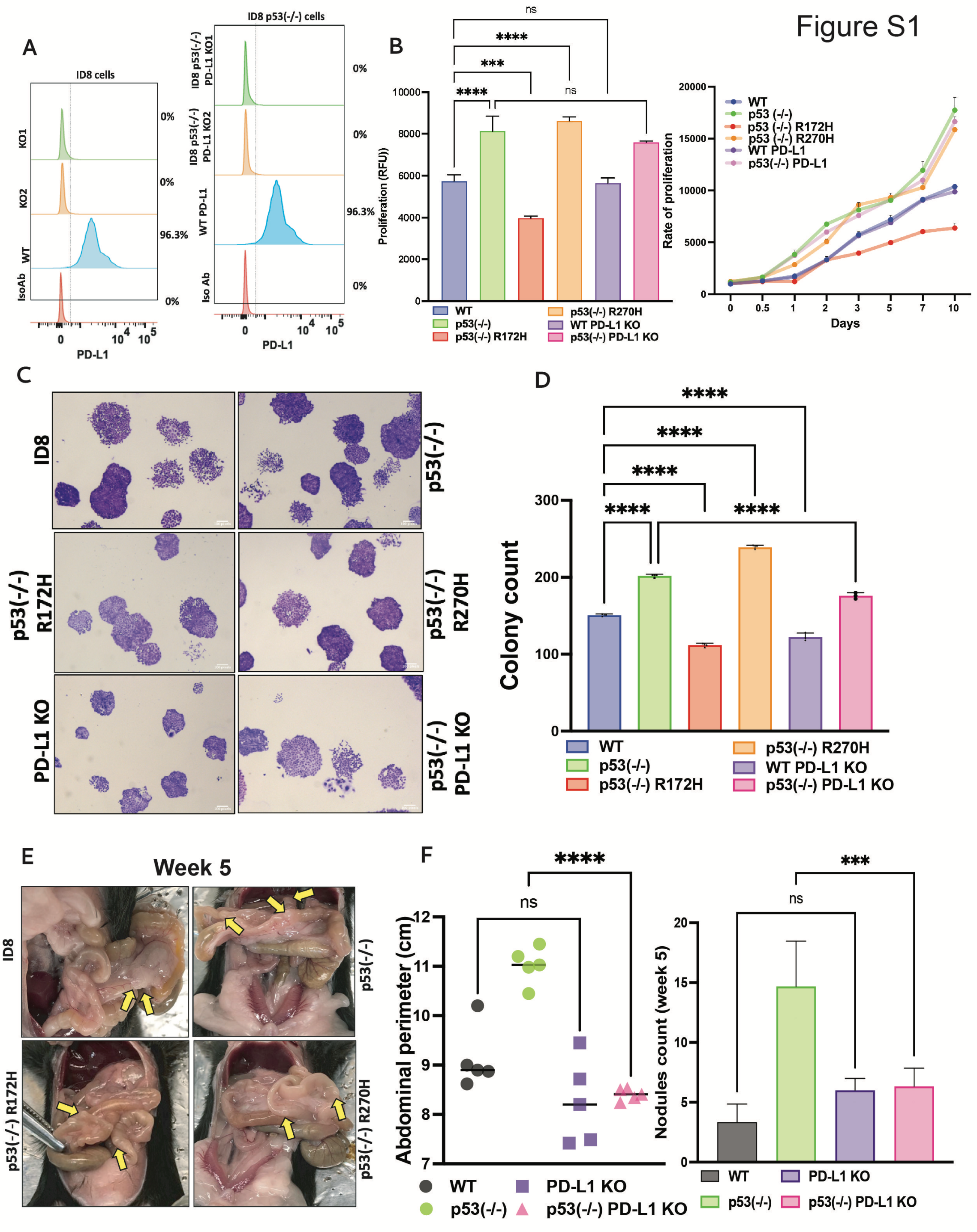


**Figure S1: Biological effects of an ectopic PD-L1 on isogenic ID8 sublines bearing p53 mutations.** (**A**) Single guide RNA (sgRNA) for the murine PD-L1 gene was designed (CTGTCTTTATATTCATGACCTAC TGG), cloned into the pX458 plasmid and then transfected into the parental ID8 and the derivative p53(-/-) cell lines. Post transfection, single GFP-positive cells were sorted in a 96-well plate and allowed to reach 80% confluence. Cell surface expression levels of PD-L1 were determined using cytometry and the indicated control cell lines. (**B**) The indicated six ID8 isogenic cell lines were seeded in 96-well plates ($3x10^3$ cells/well). After 72 hours, we assayed cell proliferation using the CellTiter Proliferation Kit (Promega). Statistical significance was assessed and the results presented as time-dependent lines or as a histogram (day 10; ns, not significant). ***, p<0.001, ****, p<0.0001 (two-tailed t-test). (**C** and **D**) The indicated cell lines were seeded in 6-well plates (500 cells/well). Media were replenished once every three days till day 14. Thereafter, cells were fixed and stained with crystal violet to count all colonies. Shown are representative images and the respective histogram (day 14). (**E**) Shown are images of the peritoneum of the indicated representative mice, which were resected 45 days post ID8 cell inoculation. Note the small, balloon-like metastatic nodules in the omentum (arrows). (**F**) Shown are histograms presenting abdominal perimeters (in cm; left panel) and metastatic nodules that were detected 5 weeks post inoculation of the indicated cells (right panel).

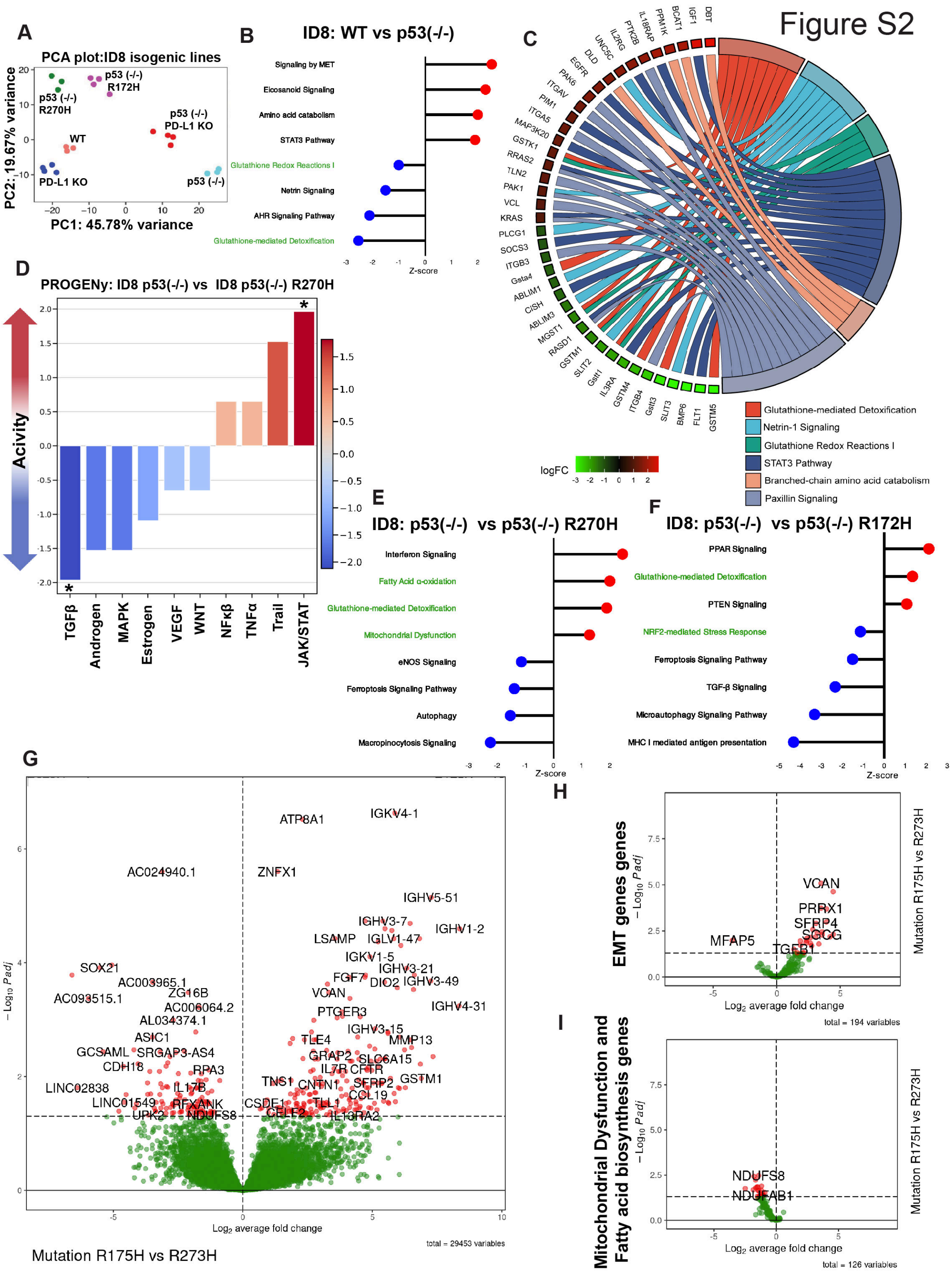


**Figure S2: Bulk RNA sequencing analysis using ID8 isogenic cell lines and an EOC patient dataset shows extensive differences between p53-R273H and p53-R175H.** (**A**) Shown are principal component analysis (PCA) clusterings of the ID8 derivative cell lines, where PC1 represents the largest variation in the transcriptomic data and PC2 represents the second largest variation. (**B**) Shown are the top significantly activated (red) and inhibited pathways (blue; absolute z-score 1 and -1, respectively) between WT cells and p53(-/-) cells. (**C**) Shown is a chord diagram, a graphical method that radially displays levels of individual transcripts. This presentation corresponds to WT versus ID8 p53(-/-) cells. (**D**) A comparative PROGENy analysis was performed on the RNA sequencing data corresponding to ID8-p53-R270H and ID8-R172H cells. These isogenic sublines are compared in terms of the activity of key signaling pathways by analyzing the expression of downstream target genes. (**E** and **F**) Shown are the top significantly activated and inhibited pathways (absolute z-score 1 and -1, respectively) comparing p53(-/-) cells versus either R172H (panel D) or R270H, both are on a null p53 background (panel E). (**G**) A volcano representation of RNA sequencing data from TCGA contrasting two p53 mutations, R273H versus R175H. Note upregulation of EMT genes (e.g., MMP13 and collagen 19) in the p53-R273H mutated tumors. (**H** and **I**) GSEA analysis was applied on the DEGs of the patient RNA-seq data comparing the R175H and R273H mutated patients' tumors showing genes responsible for EMT (panel G) or mitochondrial dysfunction and fatty acid biosynthesis (panel H) in R273H patients in comparison to R175H patients.

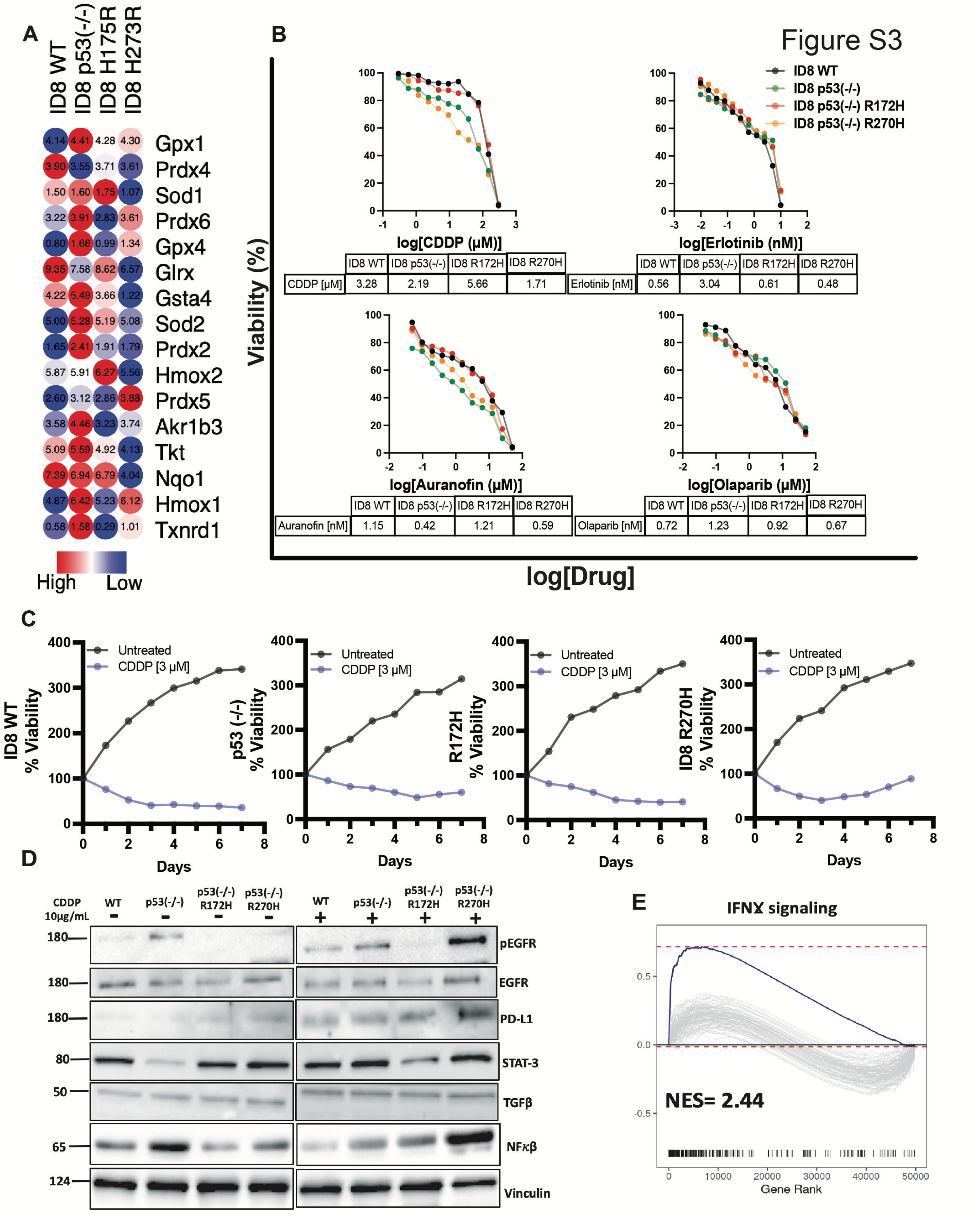


**Figure S3: The R270H mutant of p53 expressed in the ID8 isogenic line showed an increased drug sensitivity and oncogenic pathway activation relative to the p53-R172H mutant.** (**A**) qRT-PCR was performed on the indicated four isogenic cell lines to examine expression levels of specific Nrf2 targets. The results of one of three experiments are shown. (**B**) The indicated ID8 derivative cells (3x$10^3$) were seeded in 96-well plates and treated for 72 hours with Cisplatin (0-300 μM), Erlotinib (0-50 nM), Auranofin (0-50 μM) or Olaparib (0-50 μM). Cell viability was measured using the MTT (3(4,5dimethylthiazol-2yl)-2,5-diphenyltetrazolium bromide) assay. Averages of quadruplicates are shown. This experiment was repeated thrice. IC50 values are shown. (**C**) ID8 isogenic sublines were seeded in 96-well plates (500 cells/well) and treated with Cisplatin (3 μM). Media were refreshed once every three days. After nine days, cells were fixed and stained with crystal violet. The extent of cell viability determined in one of the three experiments is shown. Note that the aggressive p53 (-/-) and R270H mutant lines gained resistance to CDDP. (**D**) Four ID8 cell lines (WT, p53 null, p53-R172H, and p53-R270H) were seeded in 12-well plates at a density of $10^5$ cells per well. The cells were treated with CDDP at a concentration of 10 μg/ml per well for 24 hours. Prior to CDDP treatment, cells were subjected to a 10-hour serum starvation period. Post-treatment, cell lysates were prepared using 0.1 ml lysis buffer per well, and the lysates were analyzed using electrophoresis and immunoblotting. (**E**) Shown is a GSEA plot corresponding to differentially expressed genes in human ovarian tumors, both p53-R175H and p53-R273H (from the TCGA clinical dataset). Note upregulation of the IFN-gamma pathway in patients harboring the p53-R273H mutation.

Figure S4

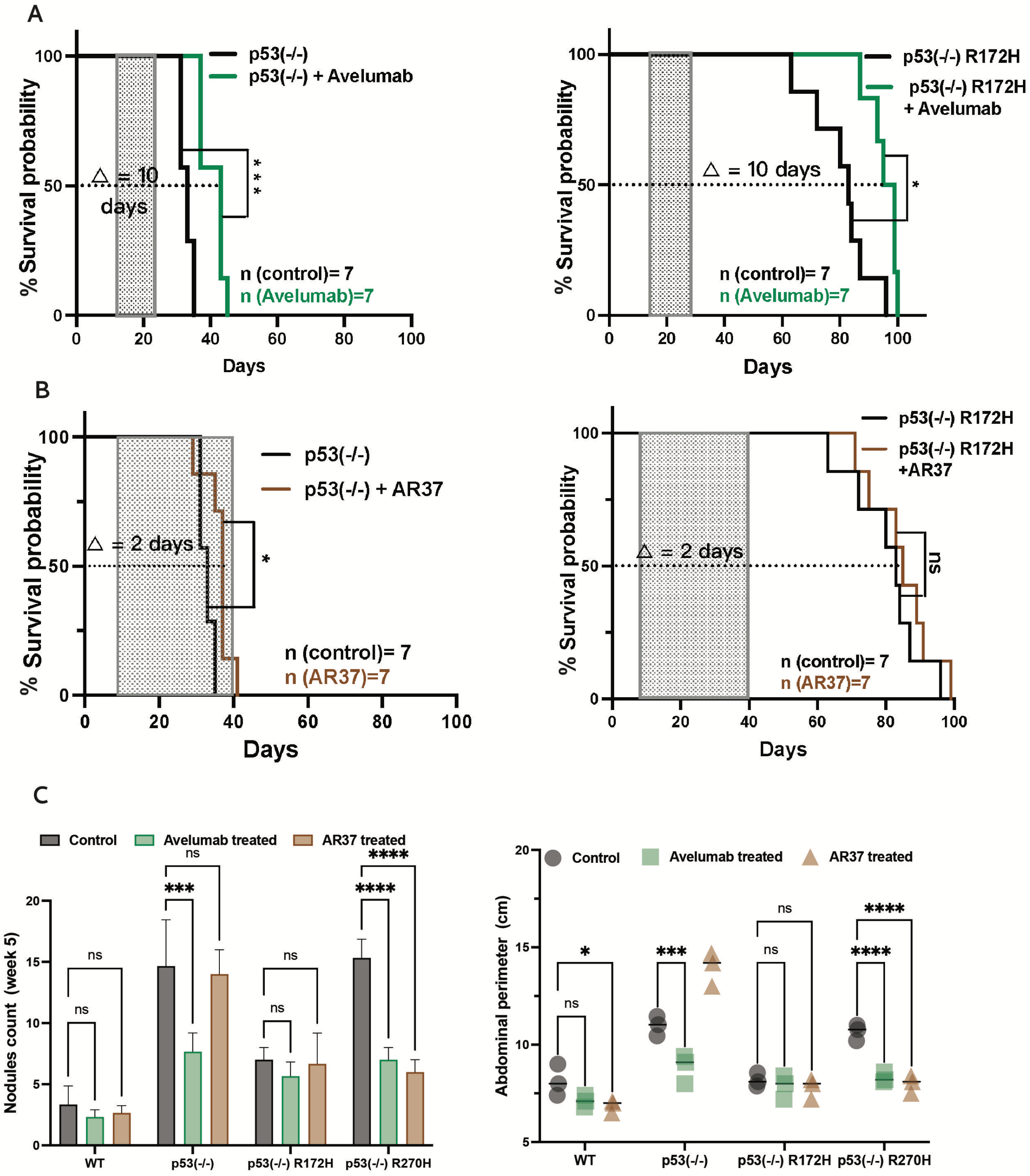


**Figure S4: The identity of p53 mutations determines the therapeutic impact of anti-AREG and, to a lesser extent, to anti-PD-L1 antibodies.** (**A**) C57/Black female mice were intraperitoneally injected with ID8 cells ($5x10^6$), either $p53^{-/-}$ (left panel) or $p53^{-/-}$ cells expressing p53-R172H (right panel). Tumor-bearing mice were randomized into two groups. One group was treated with saline, whereas the other received Avelumab (human anti-PD-L1 antibody; 0.4 mg per injection), twice weekly, starting on day ten and ending on day twenty-five. Shown are animal survival curves, along with the numbers of animals per group, survival gains and statistical parameters. (**B**) C57/Black female mice were intraperitoneally injected with ID8 cells ($5x10^6$), either $p53^{-/-}$ (left panel) or $p53^{-/-}$ cells expressing p53-R172H (right panel). Tumor-bearing mice were randomized into two groups: one group was treated with saline whereas the other was treated with AR37 (an anti-AREG antibody; 0.2

mg/injection) from day 10 to day 40. The presented survival curves include a log-rank test score and the median survival gain. *, p<0.05; n.s., non-significant. (**C**) Summary histograms presenting the number of omental nodules detected in week 10 post inoculation of cancer cells (left panel) and the respective abdominal perimeters (in cm; right panel). *, p<0.05; ***, p<0.001; ****, p<0.0001; n.s., not significant,

Figure S5

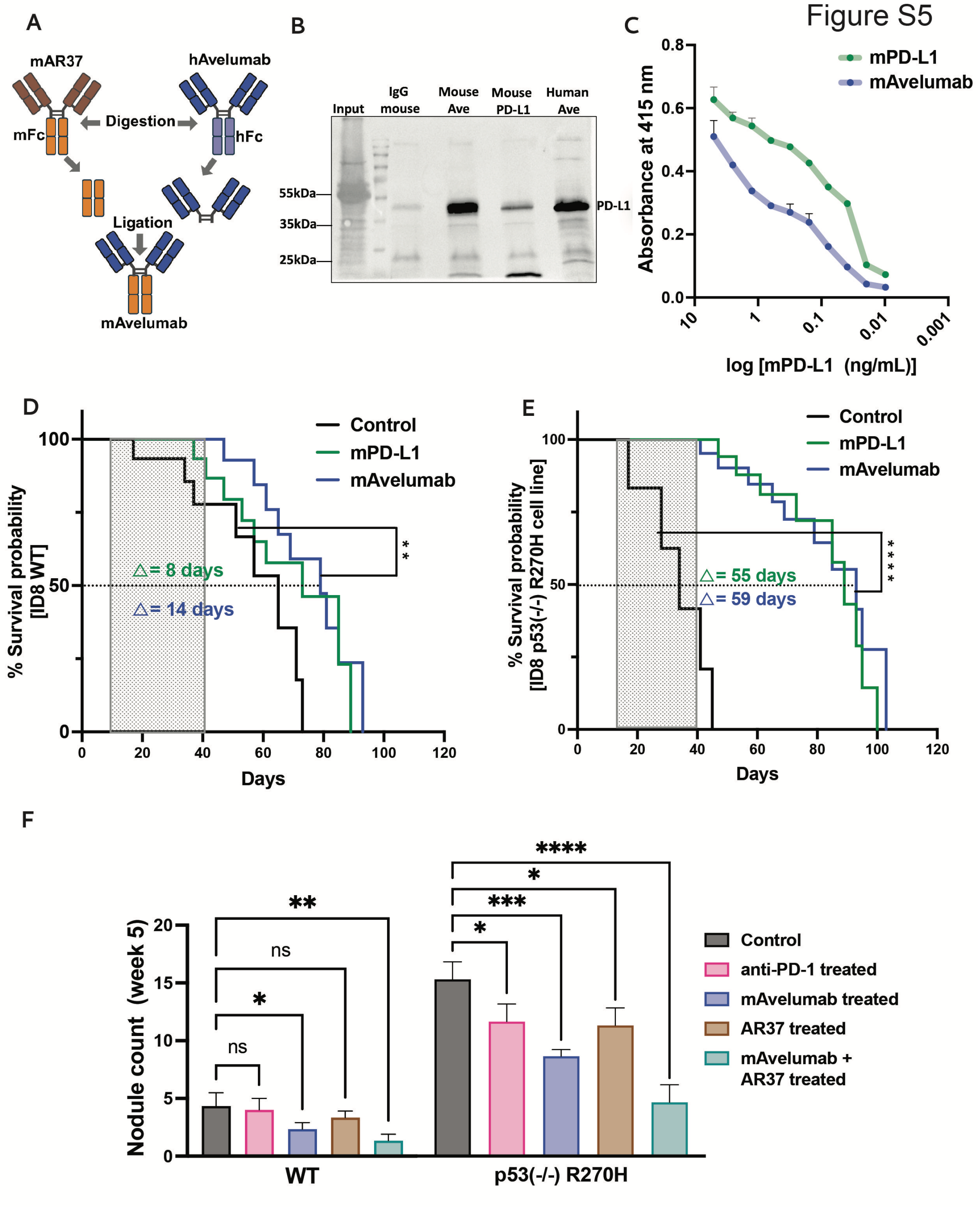


**Figure S5: Unlike the response of mice carrying p53-R270H tumors, the responses of wildtype ID8 cells to anti-PD-L1 or anti-PD-1 antibodies are relatively small.** (**A**) A schematic representation of the cloning plan of mouse avelumab outlining the replacement of Avelumab's human Fc for a murine Fc. (**B**) A recombinant mouse PD-L1 protein (extracellular domain only) was pulled down using protein

A/G agarose beads and the indicated antibodies: an engineered mouse Avelumab (murine Fc), a commercially available mouse anti-PD-L1 antibody (human Avelumab), and a control mouse immunoglobulin G (IgG). Input refers to the recombinant PD-L1 protein (1 ng). The immunoprecipitated complexes were resolved using electrophoresis and immunoblotting for PD-L1. (**C**) ELISA was performed on serial dilutions of a recombinant soluble PD-L1 that was used to coat the assay plate. Two antibodies were tested at 0.5 μg/ml: a commercial mouse anti-PD-L1 antibody (from BioXcell) and the engineered mouse Avelumab antibody. (**D** and **E**) C57/Black female mice were intraperitoneally injected with wildtype ID8 cells (D) or p53-R270H (panel E; $5x10^6$ cells). Mice were randomized 10 days later into three treatment groups (13 mice per group): untreated, commercially available mouse anti-PD-L1 antibody (0.2 mg/injection), and the self-constructed mouse Avelumab (0.2 mg/injection). All treatments were completed on day 40 post inoculation of cells (total, 10 injections). Animal survival curves are shown along with the median time gains and log-rank test scores. **, p<0.01; ****, p<0.00001 n.s., non-significant. (**F**) Quantification of metastatic nodules that emerged on the surface of the omentum of mice injected intraperitoneally with wildtype ID8 cells (left panel) or with p53-R270H cells (right panel). Nodules were counted and the results presented in histograms. Measurements were performed with 3 mice per group (in week 5). A one-way ANOVA test was performed for all of the treatment groups versus the untreated groups. *, p< 0.05, **, p< 0.01, ***, p< 0.001, ****, p<0.0001; n.s., not significant.

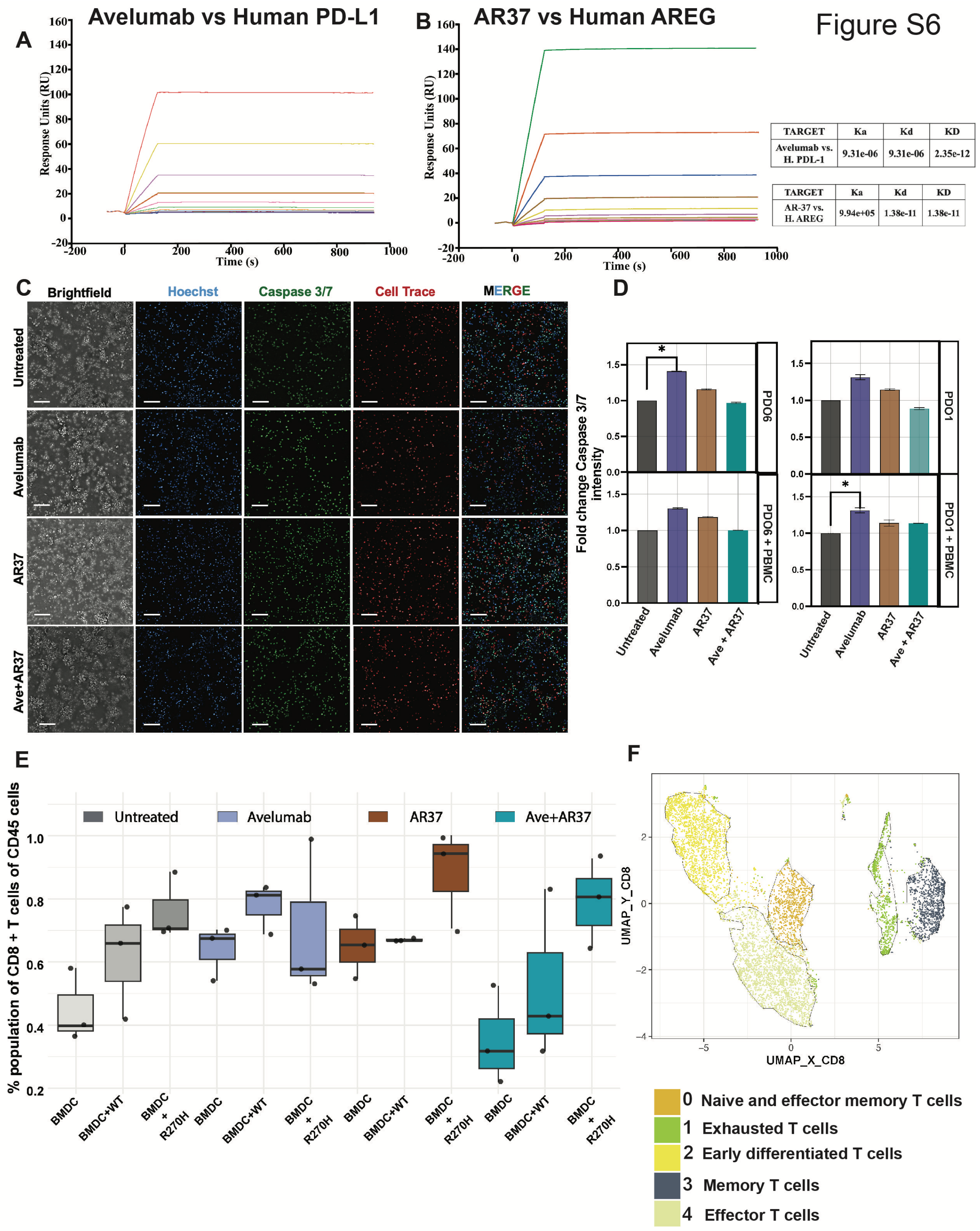


**Figure S6: Both chemo-sensitive and chemo-resistant PDO models harbouring the p53-R273C mutation respond to antibodies.** (**A**) Human PDL-1 antigen was immobilized on another channel in the same chip, and Avelumab was passed through the same channel at different concentrations. Shown are the sensograms while using the indicated antibody. (**B**) Human Amphiregulin antigen was immobilized on a channel in a chip and AR-37 was passed through the same channel at different concentrations. Shown are the sensograms while using the indicated antibody. (**C**) Shown are results of an imaging-based apoptosis assay depending on detection of caspase-3/7 activation and a fluorogenic substrate. The indicated PDOs were co-cultured with human PBMC ($3x10^5$ cells) for 5 days and treated with specific antibodies (15 micrograms per millilitre). (**D**) Quantification of the caspase activation signals only from the PDO cells is provided in histograms. Note that Avelumab displayed an apoptotic effect toward the chemo-resistant PDO6 and in the co-culture PDO1 sample. Statistical analysis used two-way ANOVA with a log-rank test score. **, $p<0.01$, ***, $p<0.001$ and ****, $p<0.0001$ is shown. Scale bar, 1.15 mm. (**E**) Cytometry-based estimation of the fraction of CD8+T cells in co-cultures comprising ID8 cells (either WT or p53(-/-) R270H) and murine BMDC, which were incubated for 2 hours in the absence or presence of the indicated antibodies. After the incubation, cells were stained for surface antigens to detect the T cell subtypes listed in Figure 6E. The CD8+ population was box-plotted across all samples. (**F**) Shown is the UMAP of the CD8+ T cells following clustering with various T cell markers, as listed in Figure 6E.

Figure S7

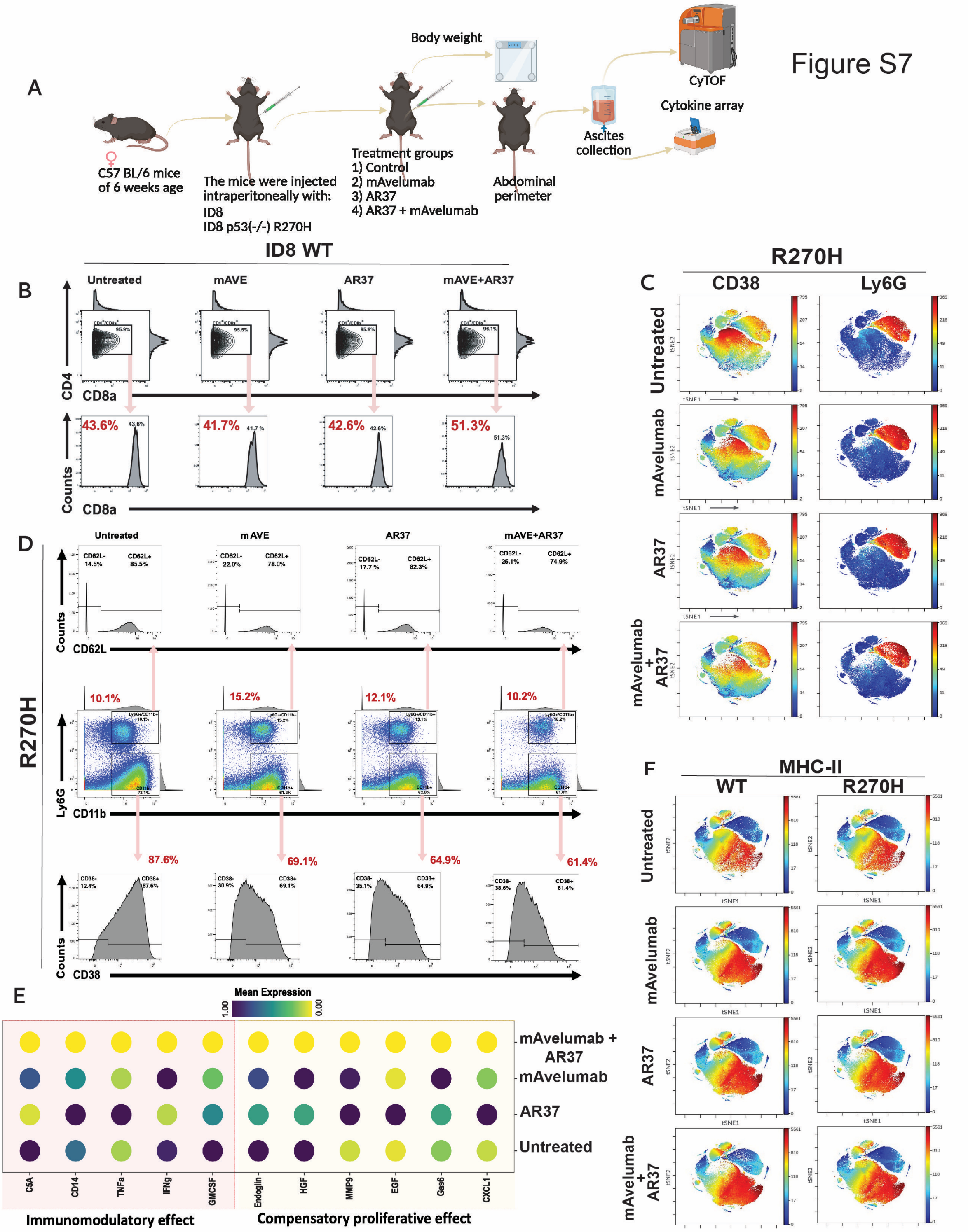


**Figure S7: Simultaneous blockade of AREG and PD-L1 boosts anti-tumor effects by means of recruiting CTLs, polarizing macrophages, diminishing neutrophils and recruiting CD38+ myeloid cells.** (**A**) A flow diagram of the experimental protocol. C57/Black female mice were intraperitoneally injected with ID8 cells (5x10$^{6}$), either WT or p53 (-/-) cells ectopically expressing p53-R270H. Ten days later, mice were randomized into four groups per cell line (5 mice per group). One group was left untreated, whereas the other groups received treatment with either mAvelumab, AR37, or a combination of mAvelumab and AR37. Ascites fluids were collected on day 75 post cell inoculation and processed for cytokine arrays and CyTOF analysis. (**B**) The CD3-positive population of the host immune cells was analyzed by applying cytometry and manual gating. This detected an increased population of CD3 and CD8 double positive T cells upon antibody combination treatment of mice bearing wildtype ID8 cells. (**C**) Using CyTOF we obtained viSNE plots reflecting decreased CD38+ myeloid cells and Lys6G+ neutrophils in the ascites fluids isolated from animals harboring p53-R270H tumors post dual antibody treatment. (**D**) Shown are analyses of the CyTOF results that made use of manual gating for CD38+ myeloid cells and Ly6G+ neutrophils. **(E)** Ascites fluids from two mice per cell line and per treatment group were isolated and overlaid on cytokine array membranes (from Proteome Profiler$^{TM}$ Mouse Cytokine Arrays). Duplicate spots corresponding to 111 cytokines were examined and data were normalized with reference to the untreated group. Relevant cytokines are presented in a dot plot showing average values of the duplicates. (**F**) Five animals per group were treated with antibodies, as indicated, and their abdominal fluids were later isolated. Shown are viSNE plots reflecting increased MHC-II antigen-presenting cells (APCs) in the fluids extracted from antibody-treated mice carrying either p53-R270H tumors or wildtype p53 tumors.
